# Sphk1-S1P signaling drives blood-brain barrier breakdown after intracerebral hemorrhage via HIF-1α-dependent upregulation of Bsg-MMP-9

**DOI:** 10.64898/2026.04.29.721777

**Authors:** Mengzhao Feng, Qi Qin, Kaiyuan Zhang, Min Yu, Fang Wang, Zhihua Li, Junlei Chang, Fuyou Guo

## Abstract

Blood-brain barrier (BBB) breakdown is a critical pathological event driving secondary brain injury and poor outcomes following intracerebral hemorrhage (ICH). However, the mechanisms governing acute BBB breakdown after ICH remain incompletely understood. Here we demonstrate that the Sphk1-S1P-S1PR3 signaling plays a pivotal role in this process. Sphk1 expression was significantly upregulated in the perihematomal endothelial cells of both ICH patients and mice, with levels positively correlating with BBB dysfunction severity and poor clinical outcomes. Using endothelial-specific genetic gain- and loss-of-function approaches, we found that Sphk1 knockdown attenuated BBB leakage, reduced hematoma volume and brain edema, preserved tight junction integrity, and improved neurological function at 1-day post-ICH, whereas Sphk1 overexpression exacerbated these pathological features. Mechanistically, transcriptomic profiling of perihematomal endothelial cells revealed that prior to its established role in Nlrp3-mediated pyroptosis, Sphk1 promotes early BBB breakdown by regulating the Bsg-MMP-9 axis. Endothelial-specific Bsg deletion completely abrogated the deleterious effects of Sphk1, confirming Bsg as an indispensable intermediary through which Sphk1 signals to MMP-9. ATAC-seq and dual-luciferase assays further demonstrated that Sphk1-generated S1P signals through S1PR3 to activate HIF-1α, which directly binds the Bsg promoter to drive its transcription, ultimately promoting MMP-9-mediated tight junction degradation. These findings delineate a complete hierarchical signaling cascade from metabolic enzyme to transcriptional regulation and subsequent barrier injury, establishing the Sphk1-Bsg-MMP-9 axis as a promising therapeutic target for ICH.

**One sentence summary:** This work identifies an early and pivotal mechanism of blood-brain barrier breakdown after intracerebral hemorrhage, demonstrating that Sphk1-generated S1P signals through S1PR3 to activate HIF-1α, which directly transactivates Bsg expression, leading to MMP-9-mediated tight junction degradation, thereby establishing a novel hierarchical axis with therapeutic potential.

## 1. Introduction

Intracerebral hemorrhage (ICH) is the most devastating subtype of stroke, associated with high mortality and long-term disability among survivors (*1, 2*). Following the primary injury caused by the hematoma, secondary brain injury driven by blood components and neuroinflammation determines clinical outcomes (*3–8*). Breakdown of the blood-brain barrier (BBB) is a central event in this process, leading to vasogenic edema, immune cell infiltration, and neuronal damage (*9–16*). However, the molecular mechanisms that initiate acute BBB breakdown after ICH remain incompletely understood.

Sphingosine kinase 1 (Sphk1) catalyzes the production of the bioactive lipid sphingosine-1-phosphate (S1P) (*17–19*). The Sphk1-S1P signaling axis has been implicated in various cerebrovascular pathologies, particularly in ischemic stroke, where it regulates endothelial barrier function and neuroinflammation through its five G protein-coupled receptors (S1PR1-5) (*20–25*). Its role in ICH, however, is less well defined. We previously reported that the Sphk1-S1P pathway contributes to BBB breakdown at 3-days post-ICH by inducing Nlrp3-mediated endothelial pyroptosis (*26*). Notably, we observed that Sphk1 inhibition as early as 1-day post-ICH, a time point preceding Nlrp3 upregulation, still conferred robust BBB protection. This finding suggested that Sphk1 may orchestrate additional, earlier-acting pathological mechanisms independent of endothelial pyroptosis.

In the present study, we sought to identify these early mechanisms and delineate the signaling cascade through which Sphk1-S1P pathway drives acute BBB breakdown following ICH. By integrating clinical samples from ICH patients, transcriptomic profiling of perihematomal endothelial cells, and complementary *in vivo* genetic approaches, we uncover a novel Sphk1-Bsg-MMP-9 signaling axis. Our findings demonstrate that Sphk1 promotes early BBB injury by transcriptionally upregulating Bsg via HIF-1α in an S1PR3-dependent manner, thereby facilitating MMP-9-mediated tight junction (TJ) degradation prior to the onset of pyroptosis. These results establish a complete hierarchical signaling pathway from metabolic enzyme to transcriptional regulation and subsequent barrier breakdown, positioning this axis as a promising therapeutic target for ICH.

## 2. Results

### 2.1 Sphk1 is upregulated in human perihematomal region after ICH and positively correlates with BBB breakdown as well as clinical severity

To investigate transcriptional alterations following ICH, we first analyzed the public human ICH dataset GSE24265. Differential expression analysis revealed 78 upregulated and 267 downregulated genes in perihematomal tissues compared to contralateral normal tissues (Fig. 1A). Among the top 20 altered genes, *SPHK1* was markedly upregulated (Fig. 1B, C). We validated this finding using clinical samples from 6 ICH patients, confirming that both *SPHK1* mRNA and Sphk1 protein levels were significantly elevated in perihematomal tissues (Fig. 1D-F). Sphk1 protein levels positively correlated with brain swelling, hematoma volume, and edema volume at admission (Fig. 1G-J), while negatively correlating with Glasgow Coma Scale (GCS) scores at admission and Glasgow Outcome Scale (GOS) scores at three months post-surgery (Fig. 1K, L). Western blotting confirmed increased extravasation of IgG and albumin in perihematomal tissues, indicating BBB breakdown (Fig. 1M-Q). Both IgG and albumin levels negatively correlated with GCS scores (Fig. 1R, S), reflecting poor clinical outcomes following BBB breakdown. Importantly, Sphk1 protein levels positively correlated with both IgG and albumin extravasation (Fig. 1T-V). These data suggest that elevated Sphk1 is associated with more severe BBB damage and worse neurological outcomes in ICH patients.

**Fig. 1.**
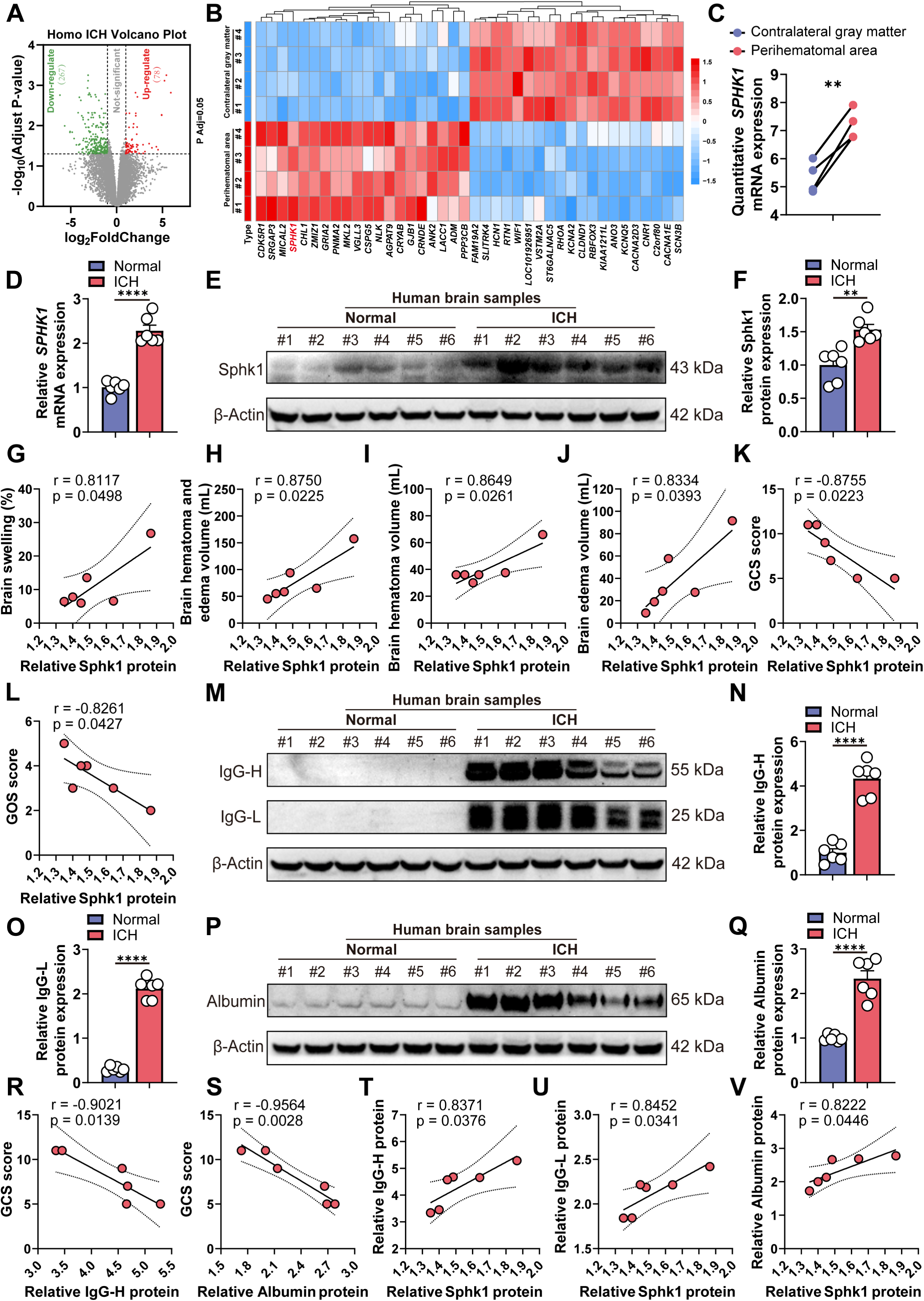
Sphk1 is upregulated in human perihematomal region after ICH and positively correlates with BBB breakdown as well as clinical severity. (A) Volcano plot of mRNA expression changes in the GSE24265 dataset after human ICH. (B) Heatmap of the top 20 up-/down-regulated genes (GSE24265). (C) Quantitative analysis of *SPHK1* expression (GSE24265). (D) RT-qPCR of *SPHK1* mRNA in normal brain tissue and perihematomal tissue from ICH patients (n=6/group). (E, F) Western blotting of Sphk1 protein in normal brain tissue and perihematomal tissue from ICH patients (n=6/group). (G-J) Correlation of Sphk1 protein expression with brain swelling, hematoma volume, and edema volume in ICH patients (n=6/group). (K, L) Correlation of Sphk1 protein expression with GCS and GOS score in ICH patients (n=6/group). (M-O) Western blotting of IgG-H and IgG-L levels in normal brain or perihematomal brain tissue from ICH patients (n=6/group). (P, Q) Western blotting of Albumin levels in normal brain or perihematomal brain tissue from ICH patients (n=6/group). (R, S) Correlation of IgG-H and Albumin with GCS score (n=6/group). (T-V) Correlation of Sphk1 expression with IgG-H, IgG-L, and Albumin (n=6/group). Data are means ± SEM; \*\**p* < 0.01, \*\*\*\**p* < 0.0001. Statistics: paired Student’s *t*-test (C); unpaired Student’s *t*-tests (D, F, N, O, Q); correlation analyses (G-L, R-V).

### 2.2 Inhibition of Sphk1 alleviates BBB breakdown and improves neurological deficits at 1 day following ICH in mice

Based on our previous finding that Sphk1 peaks within 1-day post-ICH, we administered the Sphk1 inhibitor PF543 (10 mg/kg, i.p.) at 1-hour after ICH and assessed BBB integrity at 1-day to evaluate early therapeutic efficacy in mice (Fig. 2A) (*26*). PF543 effectively suppressed ICH-induced upregulation of *Sphk1* mRNA and protein in perihematomal tissue (Fig. 2B-D). At 1-day post-ICH, PF543 significantly reduced vascular leakage, as evidenced by decreased extravasation of horseradish peroxidase (HRP), Sulfo-NHS-Biotin, and IgG compared to the ICH+Vehicle group (Fig. 2E-G). Consistently, PF543 treatment reduced hematoma volume, mitigated brain swelling on T2-weighted MRI, and attenuated midline shift (Fig. 2H, I), indicating that Sphk1 inhibition alleviates ICH-induced BBB breakdown and vasogenic edema. We next investigated the molecular basis for improved BBB function. ICH significantly downregulated TJ, effects that were reversed by PF543 treatment (Fig. 2J-M). Ultrastructural analysis by transmission electron microscopy (TEM) confirmed that PF543 reduced the proportion of open TJ between endothelial cells and decreased HRP tracer leakage into the neuropil (Fig. 2N, O). Functionally, PF543-treated mice showed progressive improvement in modified Neurological Severity Score (mNSS) from 6- to 24-hours post-ICH (Fig. 2P), enhanced motor coordination on beam walking test (Fig. 2Q), and improved sensorimotor function on adhesive tape removal test (Fig. 2R) compared to vehicle-treated ICH mice. Notably, the reduction in hematoma volume in PF543-treated mice positively correlated with improved neurological function scores (Fig. 2S-U), indicating that structural protection conferred by Sphk1 inhibition directly contributed to better functional outcomes.

**Fig. 2.**
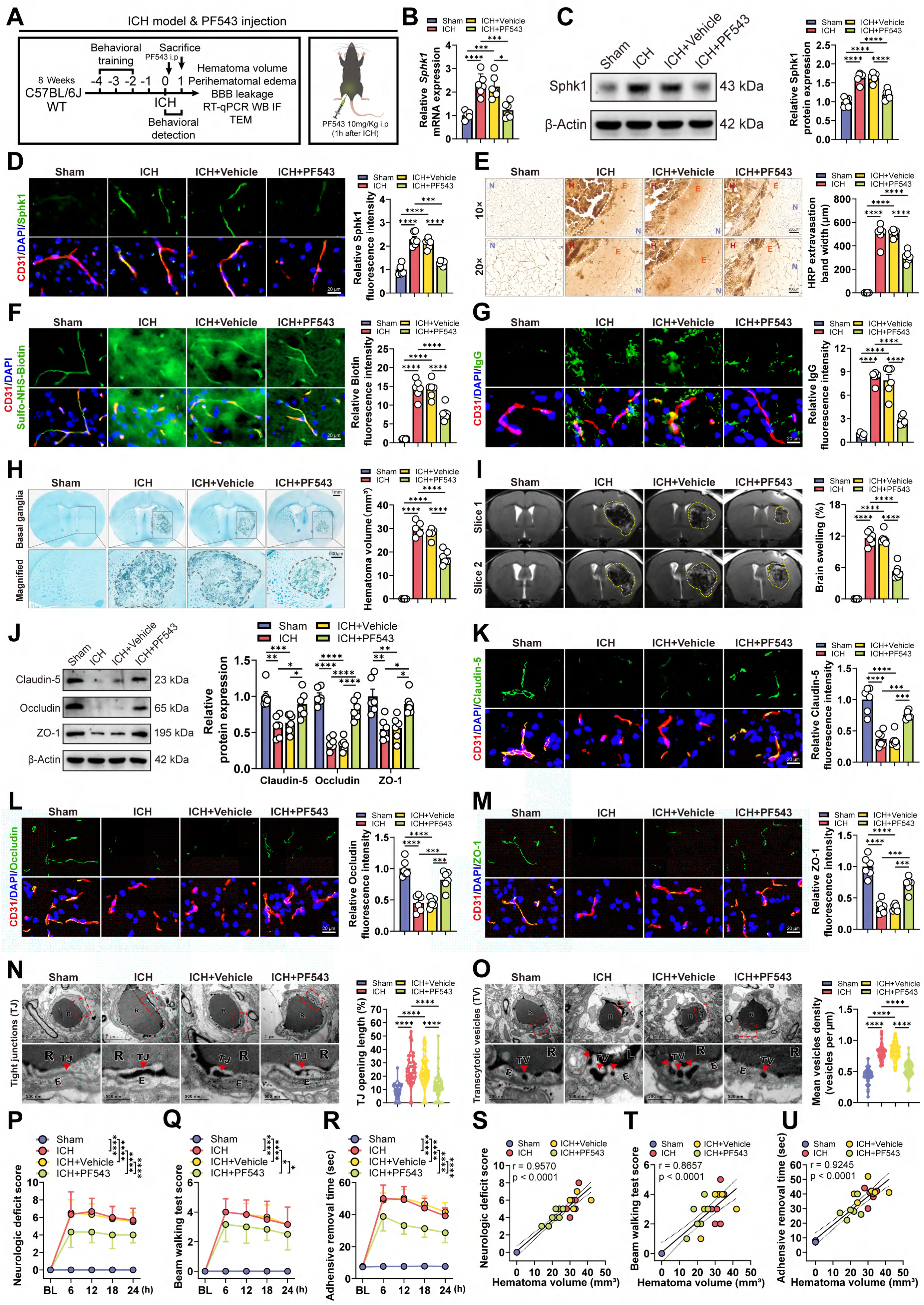
Sphk1 inhibition alleviates BBB breakdown and improves neurological deficits at 1-day following ICH in mice. (**A**) Schematic of the experimental design. (**B**) RT-qPCR analysis of *Sphk1* mRNA in the perihematomal tissue (or the corresponding brain region in Sham-operated mice) (n=6/group). (**C**) Western blotting analysis of perihematomal Sphk1 protein (n=6/group). (**D**) Immunofluorescence staining of perihematomal Sphk1 and CD31 (n=6/group). (**E**) Staining and quantification of HRP extravasation (N: Normal, E: Extravasation, H: Hematoma; n=6/group). (**F**, **G**) Immunofluorescence staining of perihematomal Sulfo-NHS-Biotin or IgG and CD31 (n=6/group). (**H**) Representative luxol fast blue staining of the hematoma (n=6/group). (**I**) Representative T2-weighted MRI showing brain swelling (n=6/group). (**J**) Western blotting analysis of perihematomal TJ proteins (n=6/group). (**K**-**M**) Immunofluorescence staining of perihematomal TJ proteins and CD31 (n=6/group). (**N**) Representative TEM images of TJ openings (arrowheads, perfused with HRP; E: Endothelial cell, R: Red blood cell) (n=6 mice/group, each data point represents one TJ). (**O**) Representative TEM of transcellular vesicles (arrowheads, perfused with HRP; TV: Transcellular vesicle) (n=6 mice/group, each data point represents one vascular field). (**P**-**R**) Neurological function assessments over time: deficit score, beam walking, and adhesive removal time (n=6/group). (**S**-**U**) Correlation analysis between hematoma volume and each neurological score (n=6/group). Data are means ± SEM; \**p* < 0.05, \*\**p* < 0.01, \*\*\**p* < 0.001, \*\*\*\**p* < 0.0001. Statistics: one-way ANOVA (**B**-**O**); two-way ANOVA with Tukey’s multiple comparisons test (**P**-**R**); correlation analyses (**S**-**U**).

### 2.3 Endothelial cell-specific knockdown of Sphk1 mitigates BBB impairment at 1-day after ICH in mice

Given our previous finding that Sphk1 upregulation after ICH is predominantly localized to brain microvascular endothelial cells, we sought to determine whether endothelial-specific modulation of Sphk1 could confer early BBB protection. Therefore, we employed a brain endothelial-targeted AAV-BI30 vector to deliver Sphk1-specific shRNA (AAV-shRNA-Sphk1) or control shRNA (AAV-shRNA-NC) via retro-orbital injection 3-weeks prior to ICH induction, and assessed BBB integrity at 1-day post-ICH in mice (Fig. 3A) (*26*). Immunofluorescence confirmed efficient and comparable transduction of brain microvascular endothelial cells in both groups (Fig. 3B). AAV-shRNA-Sphk1 significantly attenuated ICH-induced upregulation of *Sphk1* mRNA and protein in peri-hematomal tissue compared to controls, confirming successful endothelial-specific knockdown (Fig. 3C-E). Consistent with pharmacological inhibition, endothelial Sphk1 knockdown significantly reduced vascular leakage at 1-day post-ICH, as evidenced by decreased HRP and IgG extravasation (Fig. 3F-G), and led to reduced hematoma volume and alleviated brain swelling on MRI (Fig. 3H-I). Mechanistically, Sphk1 knockdown reversed ICH-induced downregulation of TJ (Fig. 3J-M), preserved TJ ultrastructure, and reduced endothelial vesicles on TEM (Fig. 3N-O). These structural improvements translated into better neurological outcomes. Mice with endothelial Sphk1 knockdown exhibited significantly lower mNSS scores, improved motor coordination on beam walking test, and enhanced sensorimotor function on adhesive tape removal test compared to controls (Fig. S1A-C). Importantly, reduced hematoma volume positively correlated with improved neurological function, indicating a direct link between structural preservation and functional recovery (Fig. S1D-F).

**Fig. 3.**
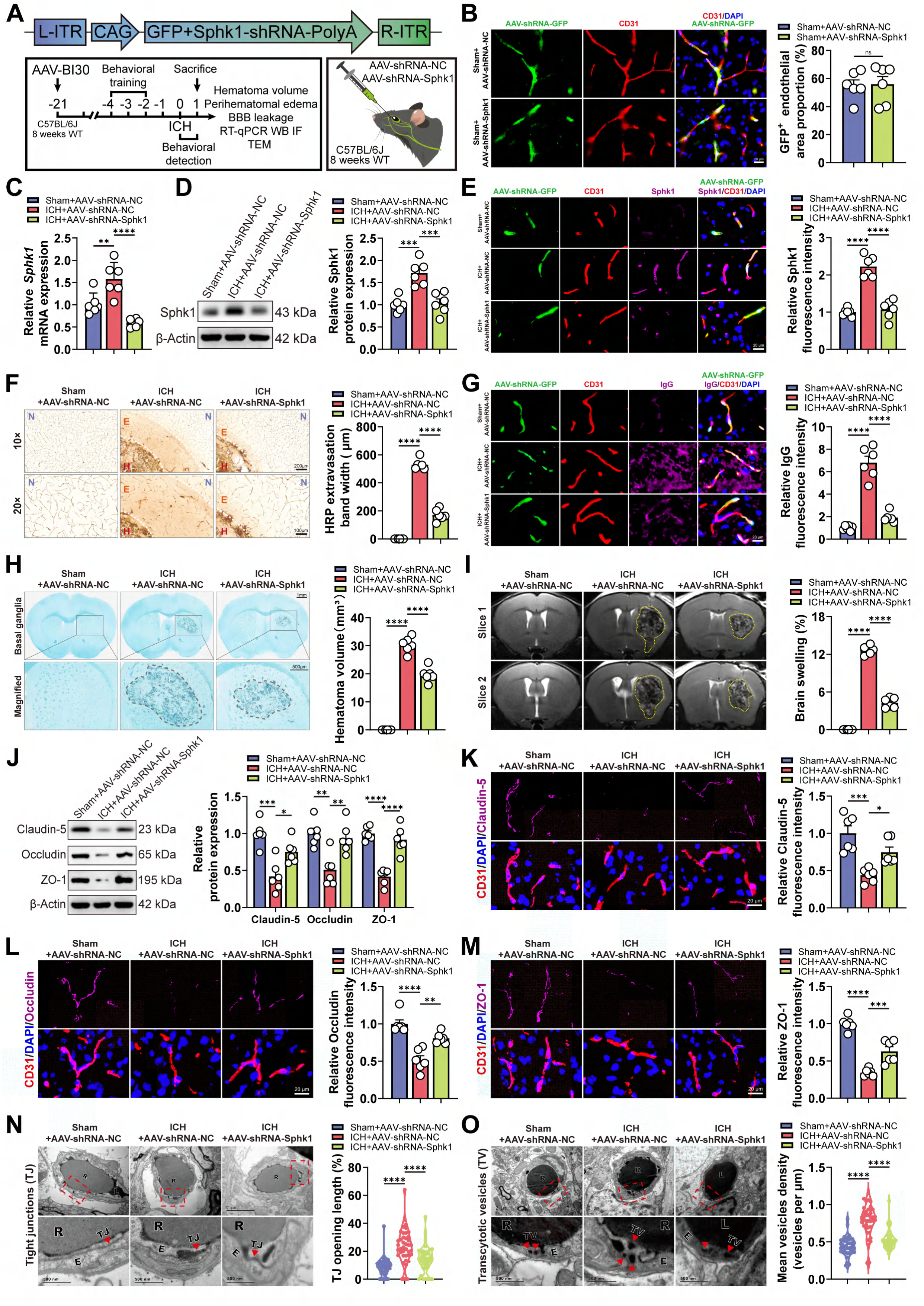
Endothelial cell-specific knockdown of Sphk1 mitigates BBB impairment at 1-day after ICH in mice. (**A**) Schematic of the experimental design. (**B**) Immunofluorescence staining of AAV-BI30-GFP and CD31 in the basal ganglia area (n=6/group). (**C**) RT-qPCR analysis of *Sphk1* mRNA in the perihematomal area (or the corresponding brain region in Sham-operated mice) (n=6/group). (**D**) Western blotting analysis of Sphk1 protein (n=6/group). (**E**) Immunofluorescence staining of GFP, CD31, and Sphk1 (n=6/group; endothelial Sphk1 signal was quantified) (**F**) Staining and quantification of HRP extravasation (N: Normal, E: Extravasation, H: Hematoma; n=6/group). (**G**) Immunofluorescence staining of IgG extravasating from endothelial cells (n=6/group). (**H**) Representative luxol fast blue staining of the hematoma in basal ganglia (n=6/group). (**I**) Representative T2-weighted MRI showing brain swelling (n=6/group). (**J**) Western blotting analysis of TJ proteins (n=6/group). (**K**-**M**) Immunofluorescence staining of TJ proteins and CD31 (n=6/group). (**N**) Representative TEM images of TJ openings (arrowheads, perfused with HRP; E: Endothelial cell, R: Red blood cell) (n=6 mice/group, each data point represents one TJ). (**O**) Representative TEM images of transcellular vesicles (arrowheads, perfused with HRP; TV: Transcellular vesicle; L: Lumen) (n=6 mice/group, each data point represents one vascular field). Data are means ± SEM; ns *p* > 0.05, \**p* < 0.05, \*\**p* < 0.01, \*\*\**p* < 0.001, \*\*\*\**p* < 0.0001. Statistics: unpaired Student’s *t*-tests (**B**); one-way ANOVA (**C**-**O**).

### 2.4 Sphk1 knockdown significantly alters the Bsg-MMP-9 mediated extracellular matrix degradation pathway

To elucidate the downstream mechanisms by which Sphk1 disrupts TJ prior to inducing endothelial pyroptosis following ICH, we performed RNA sequencing on brain microvascular endothelial cells (CD45⁻CD31⁺PDGFRβ⁻GFP⁺) isolated from the perihematomal tissue of mice at 1-day post-ICH, following prior endothelial-specific Sphk1 knockdown or control treatment (Fig. 4A). Flow cytometry confirmed successful enrichment of target endothelial populations (Fig. 4B). Comparative transcriptomic analysis identified 4907 differentially expressed genes upon Sphk1 knockdown, with 3576 downregulated and 1331 upregulated (Fig. 4C). GO and KEGG analyses confirmed effective suppression of Sphk1-S1P signaling pathways (Fig. 4D). Disease Ontology analysis revealed significant enrichment of vascular pathology-related terms among downregulated genes (Fig. 4E). Reactome pathway analysis showed pronounced downregulation of gene sets involved in extracellular matrix organization and degradation, including “Degradation of the extracellular matrix”, “Collagen degradation”, and “Activation of matrix metalloproteinases” (Fig. 4F). GSEA independently validated the negative enrichment of these ECM-related pathways and inflammatory response gene sets in the knockdown group (Fig. 4G). Core enrichment analysis consistently identified Bsg and MMP-9 as central nodes within these suppressed pathways (Fig. 4H). Their coordinated downregulation following Sphk1 knockdown nominates the Bsg-MMP-9 axis as a key downstream effector mediating Sphk1-dependent BBB breakdown at 1-day after ICH in mice.

**Fig. 4.**
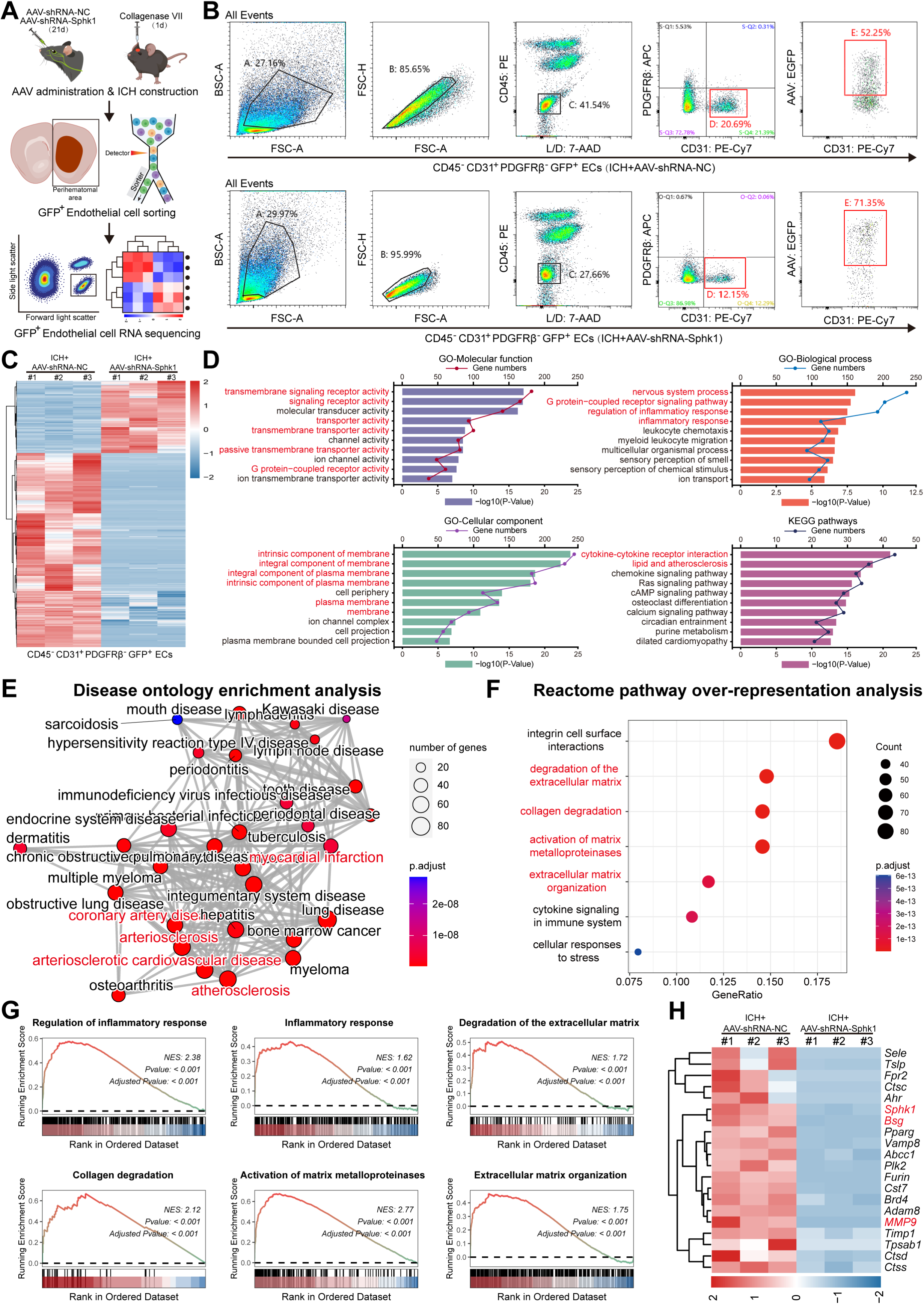
Sphk1 knockdown alters the Bsg-MMP-9-mediated extracellular matrix degradation pathway. (**A**) Schematic diagram of viral transduction, ICH modeling, tissue harvesting, and flow cytometric sorting in mice. (**B**) Gating strategy for the sorting of AAV-BI30-GFP-positive brain microvascular endothelial cells. (**C**) Heatmap of gene expression from RNA sequencing analysis. (**D**) GO and KEGG enrichment analysis for downregulated genes. (**E**) Disease ontology enrichment analysis for downregulated genes. (**F**) Reactome pathway over-representation analysis for downregulated genes. (**G**) Gene set enrichment analysis identifies key downregulated signaling pathways. (**H**) Heatmap showing expression of key downstream candidate genes.

### 2.5 Sphk1 modulates the upregulation of Bsg and MMP-9 in brain endothelial cells at 1-day after ICH in mice

To validate the Bsg-MMP-9 axis as a key transcriptional target of Sphk1, we examined its expression pattern and regulatory relationship *in vivo* (Fig. 5A). Western blotting showed that both Bsg and MMP9 protein levels were significantly increased in perihematomal tissue as early as 1-day post-ICH (Fig. 5B). Immunofluorescence co-staining revealed that upregulated Bsg specifically co-localized with the endothelial marker CD31, with minimal overlap observed for GFAP, Iba-1, or NeuN (Fig. 5C-F), identifying endothelial cells as the primary source of Bsg after ICH. In contrast, MMP-9 immunoreactivity co-localized with CD31⁺ endothelial cells, GFAP⁺ astrocytes, and Iba-1⁺ microglia, indicating a multicellular origin (Fig. 5G-J). To determine whether Sphk1 regulates this axis, we evaluated the impact of Sphk1 inhibition. Consistent with our transcriptomic data, both pharmacological inhibition with PF543 and endothelial-specific genetic knockdown of Sphk1 significantly attenuated ICH-induced upregulation of Bsg and MMP9 proteins (Fig. 5K-N).

**Fig. 5.**
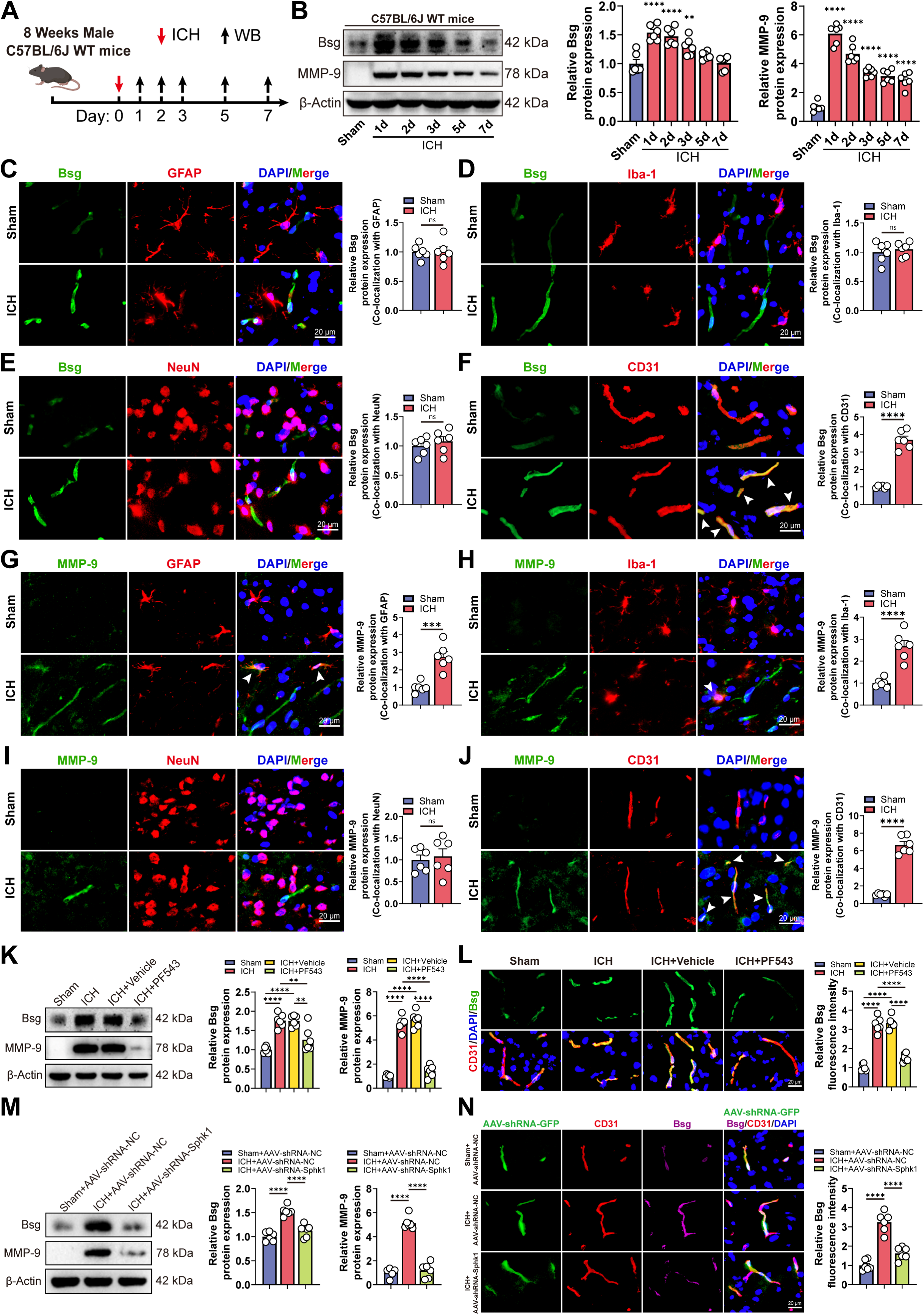
Sphk1 modulates the upregulation of Bsg-MMP-9 in vascular endothelial cells at 1-day after ICH in mice. (**A**) Experimental timeline for tissue collection and protein detection following ICH induction. (**B**) Western blotting analysis of Bsg and MMP-9 expression in perihematomal (ICH) or corresponding (Sham) brain tissue (n=6/group). (**C–F**) Immunofluorescence co-staining of Bsg with GFAP, Iba-1, NeuN, or CD31 in perihematomal or corresponding (Sham) brain tissue at 1-day after operation in mice (n=6/group). (**G–J**) Immunofluorescence co-staining of MMP-9 with GFAP, Iba-1, NeuN, or CD31 in perihematomal or corresponding (Sham) brain tissue at 1-day after operation in mice (n=6/group). (**K**) Western blotting analysis of Bsg and MMP-9 expression in the perihematomal or corresponding (Sham) brain tissue from the four indicated groups (n=6/group). (**L**) Immunofluorescence co-staining of Bsg and CD31 in the perihematomal or corresponding (Sham) brain tissue from the four indicated groups (n=6/group). (**M**) Western blotting analysis of Bsg and MMP-9 expression in the perihematomal or corresponding (Sham) brain tissue from the three indicated groups (n=6/group). (**N**) Immunofluorescence co-staining of Bsg and CD31 in the perihematomal or corresponding (Sham) brain tissue from the three indicated groups (n=6/group). Data are means ± SEM; ns *p* > 0.05, \*\**p* < 0.01, \*\*\**p* < 0.001, \*\*\*\**p* < 0.0001. Statistics: one-way ANOVA (**B**, **K**-**N**); unpaired Student’s *t*-tests (**C-J**).

### 2.6 Sphk1 overexpression promotes Bsg and MMP-9 expression and exacerbates BBB breakdown at 1-day after ICH in mice

To establish a gain-of-function link between endothelial Sphk1 and ICH pathology, we employed an AAV-BI30 vector to overexpress Sphk1 specifically in brain endothelial cells (Fig. 6A). Successful and comparable endothelial transduction was confirmed by ZsGreen fluorescence (Fig. 6B). AAV-OE-Sphk1 treatment further elevated ICH-induced *Sphk1* mRNA and protein levels (Fig. 6C-E), validating the model. Compared to controls, Sphk1 overexpression significantly aggravated BBB leakage, as shown by increased HRP and IgG extravasation (Fig. 6F, G), accompanied by larger hematoma volume, more severe brain swelling, and greater midline shift on MRI (Fig. 6H, I). Mechanistically, this was linked to further reduction of TJ and increased ultrastructural damage of endothelial junctions on TEM (Fig. 6J, K). Sphk1 overexpression also significantly amplified ICH-induced upregulation of Bsg and MMP9 proteins (Fig. 6L, M), completing the functional evidence that augmenting endothelial Sphk1 enhances the Bsg-MMP-9 axis. Consequently, mice overexpressing Sphk1 displayed significantly worse neurological outcomes on mNSS, beam walking, and adhesive tape removal tests (Fig. 6N-P), with hematoma enlargement correlating positively with neurological deficits (Fig. 6Q-S).

**Fig. 6.**
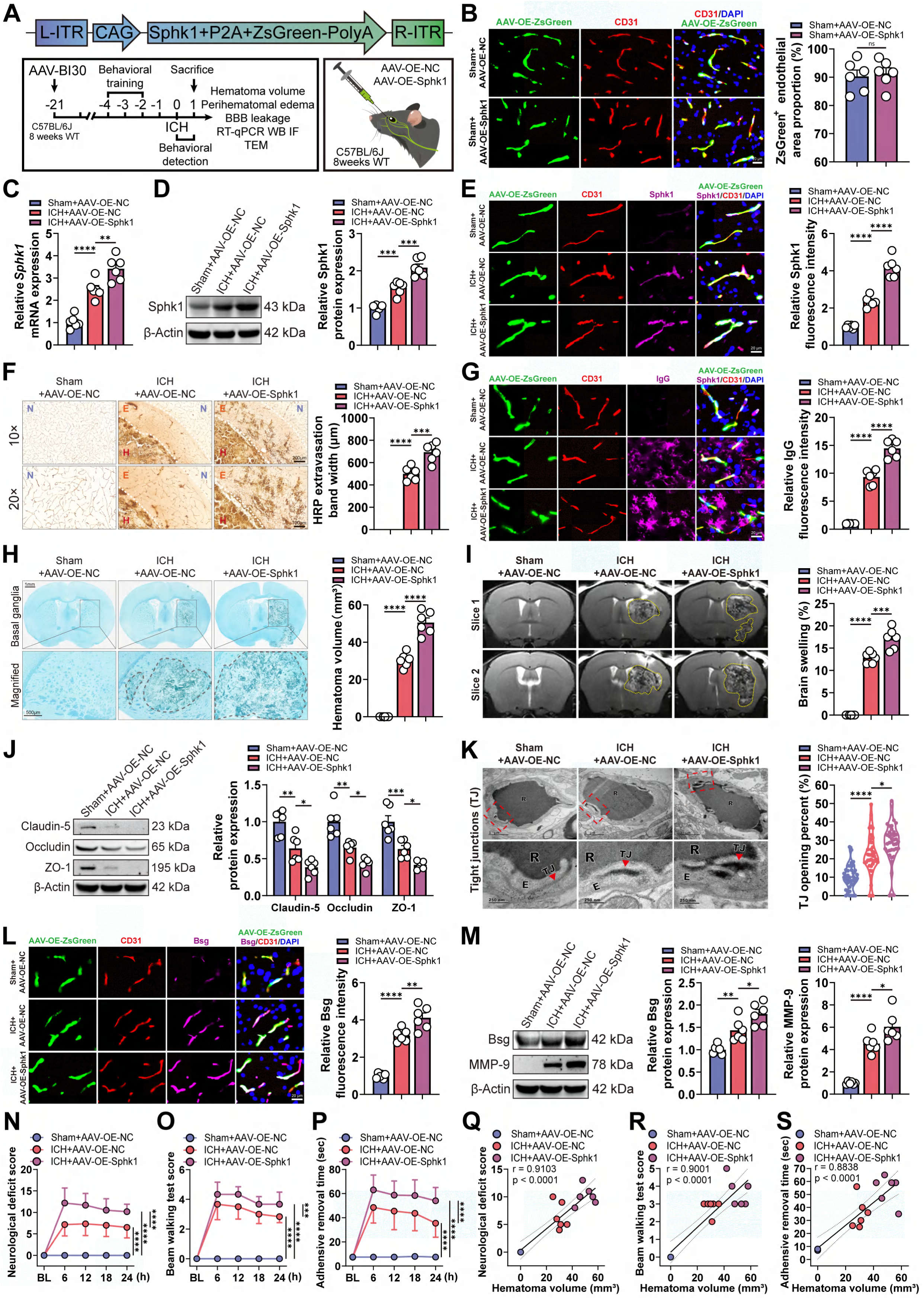
Sphk1 overexpression promotes Bsg-MMP-9 expression and exacerbates BBB breakdown at 1-day after ICH in mice. (**A**) Schematic of the experimental design. (**B**) Immunofluorescence co-staining of AAV-BI30-ZsGreen and CD31 (n=6/group). (**C**) RT-qPCR analysis of *Sphk1* mRNA in the perihematomal or corresponding (Sham) brain tissue (n=6/group). (**D**) Western blotting analysis of Sphk1 protein (n=6/group). (**E**) Immunofluorescence co-staining of Sphk1 and CD31 (n=6/group, endothelial Sphk1 signal was quantified) (**F**) Staining and quantification of HRP extravasation (n=6/group). (**G**) Immunofluorescence staining of IgG extravasation (n=6/group). (**H**) Luxol fast blue staining of hematoma (n=6/group). (**I**) T2-weighted MRI showing brain swelling (n=6/group). (**J**) Western blotting analysis of TJ proteins (n=6/group). (**K**) Representative TEM images of TJ openings (arrowheads, perfused with HRP; E: Endothelial cell, R: Red blood cell) (n=6 mice/group, each data point represents one TJ). (**L**) Immunofluorescence co-staining of Bsg and CD31 (n=6/group). (**M**) Western blotting analysis of Bsg and MMP-9 expression (n=6/group). (**N**-**P**) Neurological function assessments over time: deficit score, beam walking, and adhesive removal time (n=6/group). (**Q**-**S**) Correlation analysis between hematoma volume and each neurological score (n=6/group). Data are means ± SEM; ns *p* > 0.05, \**p* < 0.05, \*\**p* < 0.01, \*\*\**p* < 0.001, \*\*\*\**p* < 0.0001. Statistics: unpaired Student’s *t*-tests (**B**); one-way ANOVA (**C**-**M**); two-way ANOVA with Tukey’s multiple comparisons test (**N**-**P**); correlation analyses (**Q**-**S**).

### 2.7 Endothelial Bsg knockout prevents MMP-9 upregulation and alleviates BBB breakdown at 1-day after ICH in mice

To investigate the functional role of endothelial Bsg in Sphk1-mediated BBB breakdown, we generated endothelial-specific Bsg conditional knockout mice (*Bsg^flox/flox^; Cdh5-CreER^+^*, *Bsg* ECKO) and floxed littermate controls (*Bsg^flox/flox^; Cdh5-CreER^-^*, *Bsg* fl/fl) (Fig. 7A, B). Immunofluorescence confirmed significant reduction of Bsg protein in *Bsg* ECKO mice under both physiological and pathological conditions, with ICH-induced Bsg upregulation totally abrogated, confirming efficient endothelial deletion (Fig. 7C). Following ICH, *Bsg* ECKO mice exhibited significantly reduced perivascular leakage on HRP and IgG staining (Fig. 7D, E), diminished hematoma volume and brain swelling on myelin staining and MRI (Fig. 7F, G), and ameliorated downregulation of TJ proteins (Fig. 7H-K) compared to *Bsg* fl/fl controls. While Sphk1 levels were similarly elevated in both genotypes after ICH, MMP-9 upregulation was markedly blunted in *Bsg* ECKO mice, positioning Bsg downstream of Sphk1 and upstream of MMP-9 (Fig. 7L, M). Consistently, endothelial Bsg deletion significantly improved neurological outcomes, as reflected by reduced mNSS scores, better motor coordination in the beam walking test, and shorter adhesive tape removal times (Fig. 7N-P). Moreover, reduced hematoma volume correlated positively with improved neurological function, supporting a direct link between structural preservation and functional recovery (Fig. 7Q-S).

**Fig. 7.**
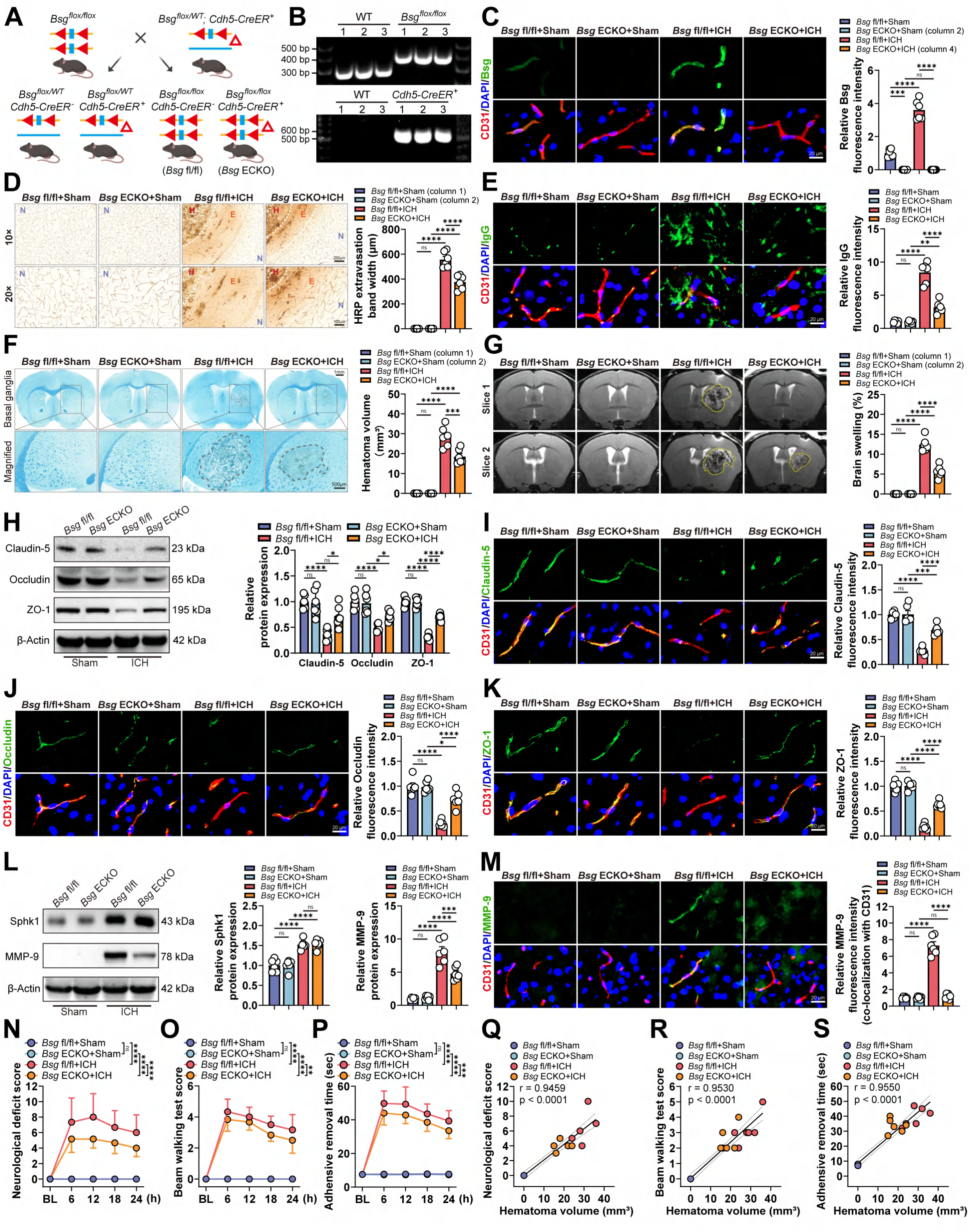
Endothelial-specific Bsg knockout prevents Sphk1-mediated MMP-9 upregulation and alleviates BBB breakdown at 1-day after ICH in mice. (**A**) Schematic of the Bsg endothelial conditional knockout mouse generation strategy. (**B**) Genotyping of *Bsg^flox/flox^* and *Cdh5-CreER^+^* alleles by agarose gel electrophoresis. (**C**) Immunofluorescence co-staining of Bsg and CD31 in perihematomal or corresponding (Sham) brain tissue (n=6/group). (**D**, **E**) BBB leakage assessed by HRP (**D**) and IgG extravasation (**E**) (n=6/group). (**F**) Luxol fast blue staining of hematoma (n=6/group). (**G**) T2-weighted MRI showing brain swelling (n=6/group). (**H**) Western blotting analysis of TJ proteins (n=6/group). (**I**-**K**) Immunofluorescence staining of TJ proteins and CD31 (n=6/group). (**L**) Western blotting analysis of Sphk1 and MMP-9 (n=6/group). (**M**) Immunofluorescence staining of MMP-9 and CD31 (n=6/group; endothelial MMP-9 signal was quantified). (**N**-**P**) Neurological function assessments over time: deficit score (**N**), beam walking (**O**), and adhesive removal time (**P**) (n=6/group). (**Q**-**S**) Correlation analysis between hematoma volume and each neurological score (n=6/group). Data are means ± SEM; ns *p* > 0.05, \**p* < 0.05, \*\**p* < 0.01, \*\*\**p* < 0.001, \*\*\*\**p* < 0.0001. Statistics: one-way ANOVA (**C**-**M**); two-way ANOVA with Tukey’s multiple comparisons test (**N**-**P**); correlation analyses (**Q**-**S**).

### 2.8 Sphk1-S1P signaling promotes HIF-1α-induced Bsg and MMP-9 expression via S1PR3

To elucidate the transcriptional mechanism by which Sphk1 regulates Bsg expression at 1-day post-ICH, we performed ATAC-seq, *in vitro* ICH modeling, and dual-luciferase reporter assays. ATAC-seq analysis revealed that endothelial Sphk1 knockdown reduced chromatin accessibility at promoter regions (12.2% vs. 14.9% in controls) and identified 6,464 downregulated peaks (Fig. 8A, B). Motif enrichment analysis revealed the bHLH domain, the core DNA-binding motif of HIF-1α, as the most significant transcription factor binding site (Fig. 8C, D). IGV visualization showed reduced chromatin accessibility at the Bsg promoter following Sphk1 knockdown (Fig. 8E), and Western blotting confirmed that ICH-induced HIF-1α upregulation was attenuated in Sphk1-knockdown mice (Fig. 8F), implicating HIF-1α as a key transcriptional mediator downstream of Sphk1. In primary brain endothelial cells subjected to hemin/hypoxia mimicking ICH, Sphk1, HIF-1α, Bsg, and MMP-9 showed time-dependent increases peaking at 1-day (Fig. 8G). PF543 treatment dose-dependently attenuated TJ downregulation (Fig. 8H) and rescued endothelial barrier function on TEER measurements (Fig. 8I), while suppressing HIF-1α, Bsg, and MMP-9 upregulation (Fig. 8J). RT-qPCR identified *S1pr3* as the most significantly upregulated S1P receptor subtype under ICH-mimetic conditions (Fig. 8K). Dual-luciferase reporter assays demonstrated that HIF-1α overexpression enhanced Bsg promoter activity, an effect significantly attenuated by either PF543 or the S1PR3 antagonist CAY10444 (Fig. 8L, M), indicating that Sphk1-generated S1P signals through S1PR3 to promote HIF-1α-mediated Bsg transcription. Collectively, these results establish that ICH activates endothelial Sphk1-S1P signaling through S1PR3 to enhance HIF-1α-mediated Bsg transcription, leading to MMP-9 upregulation and subsequent BBB breakdown at 1-day post-ICH (Fig. 8N).

**Fig. 8.**
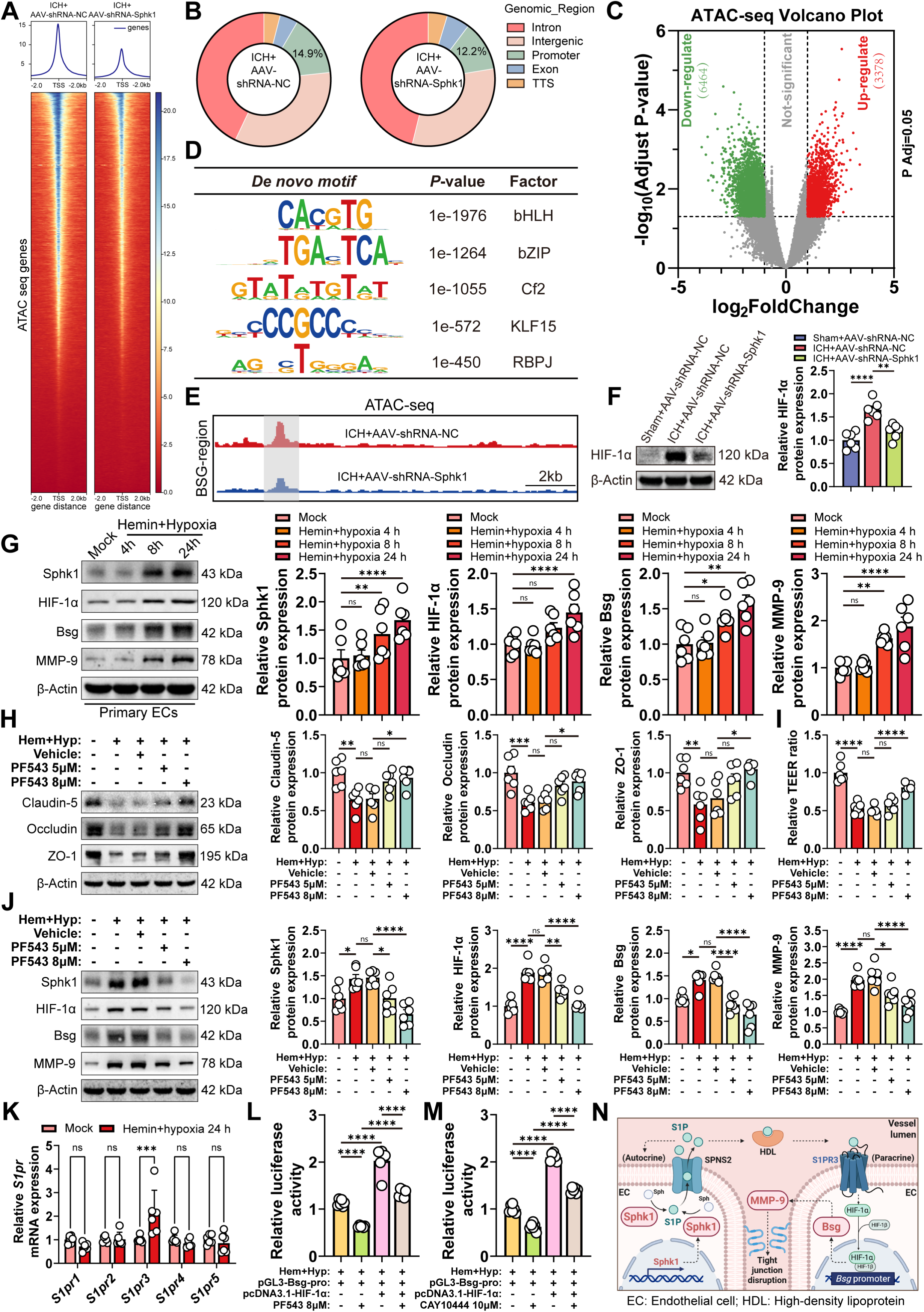
Sphk1-S1P promotes HIF-1α-induced Bsg-MMP-9 expression via S1PR3. (**A**) ATAC-seq analysis of chromatin accessibility in perihematomal tissue from ICH mice with Sphk1 knockdown or control (n=3/group). (**B**-**E**) Bioinformatics analysis including annotation (**B**), volcano plot (**C**), motif enrichment (**D**), and IGV tracks at the *Bsg* promoter (**E**). (**F**) Western blotting of perihematomal HIF-1α after ICH with or without Sphk1 knockdown in mice (n=6/group). (**G**) Western blotting of Sphk1, Bsg, HIF-1α, and MMP-9 under ICH-mimicking conditions in primary mouse brain endothelial cells (n=6/group). (**H**) Western blotting of TJ in primary mouse brain endothelial cells under ICH-mimicking conditions with or without Sphk1 inhibition (n=6/group). (**I**) TEER measurements in primary mouse brain endothelial cells under ICH-mimicking conditions with or without Sphk1 inhibition (n=6/group). (**J**) Western blotting of Sphk1, HIF-1α, Bsg, and MMP-9 in primary mouse brain endothelial cells under ICH-mimicking conditions with or without Sphk1 inhibition (n=6/group). (**K**) RT-qPCR of *S1pr1-5* in primary mouse brain endothelial cells under ICH-mimicking conditions (n=6/group). (**L**, **M**) Luciferase reporter assay for HIF-1α activity on the *Bsg* promoter in HEK293T cells under ICH-mimicking conditions with Sphk1 or S1PR3 inhibition (n=6/group). (**N**) Schematic diagram. Data are means ± SEM; \**p* < 0.05, \*\**p* < 0.01, \*\*\**p* < 0.001, \*\*\*\**p* < 0.0001. Statistics: one-way ANOVA (**F**-**J**, **L**, **M**); two-way ANOVA with Tukey’s test (**K**).

## 3. Discussion

The present study identifies a novel molecular axis, Sphk1-Bsg-MMP-9, as a critical driver of BBB breakdown following ICH. We provide multi-layered evidence for this cascade. Sphk1 was significantly upregulated in peri-hematomal endothelial cells of both ICH patients and mice, with levels correlating directly with BBB compromise and poor neurological outcomes. Pharmacological inhibition or endothelial-specific knockdown of Sphk1 preserved BBB integrity, reduced cerebral edema, and improved neurobehavioral function, effects linked to restoration of TJ and suppression of basement membrane degradation. Conversely, endothelial Sphk1 overexpression exacerbated injury. Transcriptomic analysis of Sphk1-deficient endothelial cells identified the Bsg-MMP-9 axis as a key downstream effector, a finding validated by gain- and loss-of-function experiments. Using endothelial-specific Bsg conditional knockout mice, we confirmed Bsg as an indispensable intermediary, as its deletion reversed the deleterious effects of Sphk1. Mechanistically, we demonstrate that Sphk1-generated S1P signals through S1PR3 to stabilize and activate HIF-1α, which directly transactivates the Bsg promoter, initiating MMP-9-mediated BBB damage.

The BBB is a crucial neurovascular structure that maintains brain homeostasis, and its breakdown is a key pathological event in various neurological diseases (*27–38*). It has been identified as a pivotal trigger for secondary brain injury and poor outcomes following ICH (*39–41*). We previously reported that the Sphk1-S1P axis contributes to BBB damage at 3-days post-ICH by inducing Nlrp3-mediated endothelial pyroptosis (*26*). The present study was motivated by the observation that Sphk1 inhibition conferred significant cerebrovascular protection as early as 1-day post-ICH, a time point preceding the peak of Nlrp3-driven pyroptosis. This temporal discrepancy suggested that Sphk1 orchestrates a broader assault on the BBB through multiple complementary pathways acting in sequence. The current work substantiates this hypothesis by elucidating a novel early mechanism, showing that prior to the onset of pyroptosis, the Sphk1-S1P axis promotes inter-endothelial TJ degradation through transcriptional upregulation of the Bsg-MMP-9 cascade. This early breakdown likely sets the stage for subsequent pyroptotic injury by increasing the vulnerability of the neurovascular unit.

At 1-day post-ICH, the mechanisms by which Sphk1 inhibition or endothelial-specific knockdown confers BBB protection remained unexplored. While our previous work showed that continuous Sphk1 inhibition for 3-days attenuated BBB injury by suppressing Nlrp3-mediated pyroptosis, we observed that a single PF543 administration at 1-hour post-ICH, or prior endothelial-specific Sphk1 knockdown, was sufficient to elicit marked BBB protection at 1-day, a time point when Nlrp3 expression was not yet elevated (*26*). This temporal dissociation suggested that Sphk1 operates through additional earlier-acting pathways independent of pyroptosis. To investigate this hypothesis, we performed transcriptomic profiling on FACS-purified brain microvascular endothelial cells from the perihematomal region at 1-day post-ICH, specifically selecting those transduced with AAV-BI30-shRNA targeting Sphk1 or control vectors. This unbiased approach identified the Bsg-MMP-9 axis as a prominent downstream pathway modulated by Sphk1 during acute ICH. We validated these findings at the protein level, demonstrating that pharmacological inhibition or genetic knockdown of Sphk1 suppressed ICH-induced upregulation of Bsg and MMP-9, while endothelial-specific Sphk1 overexpression potentiated their expression and exacerbated BBB injury. These complementary experiments demonstrate that within the acute 1-day window following ICH, Sphk1 drives BBB breakdown by transcriptionally regulating the Bsg-MMP-9 cascade, promoting early TJ degradation independently of later pyroptotic cell death. This positions Sphk1 as a central early orchestrator of endothelial dysfunction, initiating a coordinated pathological sequence that sensitizes the neurovascular unit for subsequent inflammatory injury.

The Bsg-MMP-9 axis has been implicated in secondary brain injury following ICH, with Bsg serving as an inducer of MMP-9 that contributes to BBB breakdown through degradation of TJ and basement membrane components (*42–44*). Although Bsg-MMP-9 is recognized as a critical effector pathway in ICH pathology, the upstream mechanisms governing its activation have remained largely unexplored. The present study addresses this knowledge gap by identifying the Sphk1-S1P signaling cascade as a novel upstream regulator of the Bsg-MMP-9 axis. Using complementary approaches including pharmacological inhibition, endothelial-specific gene knockdown, and overexpression, we demonstrate that under pathological ICH conditions, the expression of both Bsg and MMP-9 is dynamically modulated by Sphk1 activity. To determine whether Bsg serves as an indispensable intermediary linking Sphk1 to MMP-9-mediated BBB injury, we generated an endothelial-specific Bsg conditional knockout mouse line (*Bsg^flox/flox^*; *Cdh5-CreER^+^*). In this genetic model, endothelial-specific overexpression of Sphk1 followed by ICH induction revealed that Bsg deficiency profoundly abrogated the exacerbated BBB pathology associated with Sphk1. The protective effects of Bsg deletion were observed despite sustained elevation of Sphk1, confirming that Bsg functions as an essential signaling relay downstream of Sphk1 and upstream of MMP-9. These rescue experiments provide definitive genetic evidence establishing Bsg as a necessary intermediary in the Sphk1-driven cascade leading to MMP-9 activation and subsequent BBB breakdown.

Having established Sphk1 as an upstream regulator of the Bsg-MMP-9 axis, we next sought to elucidate the transcriptional mechanism by which Sphk1-S1P signaling governs Bsg expression. ATAC-seq analysis of endothelial cells with Sphk1 knockdown revealed a significant reduction in chromatin accessibility at the Bsg gene promoter following ICH, indicating transcriptional regulation by Sphk1-S1P signaling. Motif analysis of the downregulated peaks identified the bHLH motif as the most significantly enriched transcription factor binding site. The bHLH domain is the critical DNA-binding component of HIF-1α, which confers target gene specificity to the HIF-1 transcriptional complex (*45–47*). This chromatin-based finding pointed to HIF-1α as a prime mediator of Sphk1-driven Bsg transcription. Consistent with this hypothesis, previous studies have demonstrated that S1P can activate HIF-1α in vascular endothelial cells under physiological conditions (*48*). Our study significantly extends this concept by demonstrating, for the first time under the pathological condition of ICH, that the Sphk1-S1P axis actively regulates HIF-1α expression in cerebral endothelial cells. To validate our sequencing findings, we performed dual-luciferase reporter assays under conditions mimicking the ICH microenvironment, confirming that HIF-1α binds directly to the Bsg promoter to drive its transcriptional activity. This aligns with reports in epithelial solid tumors, where hypoxia-induced HIF-1α binding to the Bsg promoter promotes tumor progression (*49*). The present study extends these findings by demonstrating that within the unique microenvironment of ICH, the Sphk1-S1P signaling pathway promotes HIF-1α binding to the Bsg promoter, thereby upregulating Bsg expression and exacerbating BBB injury.

To elucidate the receptor-mediated mechanism by which Sphk1-generated S1P initiates HIF-1α activation, we examined the expression of individual S1P receptor subtypes following ICH. Among the five known receptors (S1PR1-5), S1PR3 exhibited the most pronounced upregulation in response to ICH pathology. This observation aligns with and significantly extends our previous work demonstrating that S1PR3 contributes to BBB breakdown and neuroinflammation after ICH, and that its inhibition with CAY10444 attenuates these pathological processes (*50*). In the present study, we confirmed that S1PR3 is the predominantly upregulated receptor subtype. Critically, dual-luciferase reporter assays revealed that pharmacological antagonism of S1PR3 with CAY10444 significantly reduced HIF-1α binding to the Bsg promoter, thereby attenuating its transcriptional activity. These findings demonstrate that the Sphk1-S1P axis transduces the pathological ICH stimulus into the intracellular compartment specifically through S1PR3, which subsequently promotes HIF-1α-mediated transcription of Bsg and downstream induction of MMP-9.

The present study acknowledges certain limitations. While our findings provide robust preclinical evidence supporting Sphk1 as a therapeutic target, clinical translation has not yet been achieved. Future development of Sphk1-targeted interventions for ICH patients will require rigorous evaluation of safety, efficacy, and optimal therapeutic windows in appropriate large animal models, followed by well-designed clinical trials. Additionally, although the collagenase-induced ICH model employed herein is widely accepted and recapitulates key features of human ICH, it may not fully capture the heterogeneity and complexity of spontaneous ICH. Future studies utilizing alternative models, such as autologous blood injection, would further validate the generalizability of our findings.

In conclusion, this study identifies the Sphk1-Bsg-MMP-9 signaling axis as a critical driver of BBB breakdown following ICH. We establish clinical relevance by correlating Sphk1 expression with patient outcomes, confirm causality through endothelial-specific genetic manipulation, and elucidate a complete hierarchical mechanism wherein Sphk1-generated S1P signals via S1PR3 to activate HIF-1α-mediated transcription of Bsg, promoting MMP-9-dependent TJ degradation. Notably, Sphk1 orchestrates a temporally coordinated, multi-faceted injury to the neurovascular unit, promoting early Bsg-MMP-9-mediated barrier breakdown prior to its later role in Nlrp3-driven endothelial pyroptosis, with Bsg serving as an indispensable intermediary. Collectively, these findings position this novel signaling pathway as a promising multi-target therapeutic strategy for ICH.

## 4. Materials and methods

### Bioinformatics analysis of GEO dataset

The gene expression dataset GSE24265 was downloaded from the Gene Expression Omnibus (GEO; https://www.ncbi.nlm.nih.gov/geo/). This dataset contains transcriptomic profiles of perihematomal tissue, contralateral grey matter, and contralateral white matter from four patients who died of supratentorial ICH. Differentially expressed genes between perihematomal and contralateral grey matter were identified using the limma package in R with thresholds of adjusted *p* < 0.05 and ∣log_2_(fold change)∣> 0.5. Results were visualized by volcano plots and heatmaps.

### Human brain tissue samples

In patients with acute spontaneous basal ganglia ICH who underwent emergency surgery at the First Affiliated Hospital of Zhengzhou University, perihematomal brain tissue and surgically accessible normal brain tissue that was necessarily removed during the surgical approach to expose the hematoma were collected from six patients. All samples were snap-frozen in liquid nitrogen and stored at −80 °C. The study was approved by the hospital ethics committee (approval No. 2021-KY-0156) and written informed consent was obtained from authorized families of patients. Admission GCS scores and GOS score at three months after surgery were recorded.

### Animals and ICH model

Adult male C57BL/6J mice (8-10 weeks old, 20-25 g) were obtained from Beijing Vital River Laboratory Animal Technology and housed under specific pathogen-free conditions with a 12-hours light/dark cycle. ICH was induced by stereotaxic injection of 0.15 U collagenase VII-S (C2399, Sigma-Aldrich) dissolved in 0.5 μL sterile saline into the right striatum (coordinates: 0.3 mm anterior, 2.3 mm lateral, 3.0 mm ventral to bregma). Sham-operated mice received saline only. All animal procedures were approved by the Institutional Animal Care and Use Committee of Shenzhen Institutes of Advanced Technology, Chinese Academy of Sciences (approval No. SIAT-IACUC-231116-FMZ-A2385).

### AAV-mediated gene manipulation

To achieve brain endothelial-specific knockdown of Sphk1, an adeno-associated virus (AAV-BI30) (*51*) carrying a short hairpin RNA against mouse *Sphk1* (AAV-shRNA-Sphk1) or a scrambled control (AAV-shRNA-NC) was constructed (WeiZhen Biotech). For overexpression, AAV-BI30 carrying mouse *Sphk1* coding sequence (AAV-OE-Sphk1) or empty vector (AAV-OE-NC) was generated. Viruses (5×10^11^ vg per mouse) were administered via retro-orbital injection 3-weeks before ICH induction.

### Endothelial-specific Bsg knockout mice

Bsg-flox mice (*Bsg^fl/fl^*) were generated by CRISPR/Cas9 technology (C57BL/6JGpt-*Bsg^em1Cflox^*/Gpt, GemPharmatech, Strain NO. T057905) and crossed with *Cdh5-CreER* mice (*52*) (generously provided by Ralf Adam, Max Plank Institute of Molecular Biomedicine, Germany). 8-10-weeks male *Bsg^fl/fl^; Cdh5-CreER^+^* (*Bsg* ECKO) and *Bsg^fl/fl^; Cdh5-CreER^−^* (*Bsg* fl/fl) littermates received intraperitoneal tamoxifen (75 mg/kg, 10540-29-1, Sigma-Aldrich) every other day for four doses, followed by a 14-day washout period before ICH surgery.

### Primary brain microvascular endothelial cell culture

Primary mouse brain microvascular endothelial cells were isolated from 8-10-weeks C57BL/6J mice as previously described (*28*). Briefly, cerebral cortices were minced and digested in Hanks’ balanced salt solution containing 1 mg/mL neutral protease, 1 mg/mL collagenase type 4, and 20 U/mL DNase I at 37 °C for 45-minutes. After filtration through a 40-μm cell strainer, the cell suspension was centrifuged in 20% BSA at 1000× g for 25 min to remove myelin. The pellet was resuspended in endothelial growth medium (ECM, CM-ZY001, Procell) supplemented with 4 μg/mL puromycin and cultured at 37 °C with 5% CO₂. To mimic the ICH microenvironment *in vitro*, primary brain endothelial cells were exposed to 1 μM hemin (51280, Sigma-Aldrich) and placed in a hypoxia chamber (Billups-Rothenberg) flushed with 95% N₂/5% CO₂ for 24-hours.

### RT-qPCR

Total RNA was extracted from human or mouse tissues using the FastPure Cell/Tissue Total RNA Isolation Kit (Vazyme). Reverse transcription was performed with HiScript III RT SuperMix (R232-01, Vazyme). Quantitative PCR was carried out using ChamQ SYBR qPCR Master Mix (Q341-02, Vazyme) on a LightCycler 96 system (Roche). Relative mRNA levels were calculated by the 2^−ΔΔCt^ method with ACTB or Actb as internal control. The following primer sequences were used: Human: *SPHK1* forward primer: TCTGCTTGGTCCAATGTGCAA, *SPHK1* reverse primer: GGAACAGTTCGTGTCATCCTC. *ACTB* forward primer: GCTGCATTTAGTGGCCTCATT, *ACTB* reverse primer: GCAAGGCATAACCTGATGTGG. Mouse: *Sphk1* forward primer: GCAACGTGGAATCACCACTGA, *Sphk1* reverse primer: CAGCCAGTAGTCTGTGGACTC. *S1pr1* forward primer: ATGGTGTCCACTAGCATCCC, *S1pr1* reverse primer: TGCCTGTGTAGTTGTAATGCC. *S1pr2*forward primer: ACAGCAAGTTCCACTCAGCAA, *S1pr2* reverse primer: CTGCACGGGAGTTAAGGACAG. *S1pr3* forward primer: ACTCTCCGGGAACATTACGAT, *S1pr3* reverse primer: CCAAGACGATGAAGCTACAGG. *S1pr4* forward primer: CTGGCTACTGGCAGCTATCC, *S1pr4* reverse primer: AGACCACCACACAAAAGAGCA. *S1pr5* forward primer: CCTGCTTCGTACCCTTAGCG, *S1pr5* reverse primer: GGCACGCGACATCCAGTAAT. *Actb* forward primer: ATGACCCAAGCCGAGAAGG, *Actb* reverse primer: CGGCCAAGTCTTAGAGTTGTTG.

### Western blotting

Proteins were extracted from tissues or cells using RIPA buffer containing protease and phosphatase inhibitors (4906845001, Roche). Equal amounts of protein were separated by SDS-PAGE, transferred to PVDF membranes, blocked with 5% non-fat milk, and incubated overnight at 4 °C with primary antibodies against Sphk1 (1:1000, 10670-1-AP, Proteintech), Albumin (1:1000, 16475-1, Proteintech), Bsg (1:1000, ab188190, Abcam), MMP-9 (1:1000, ab283575, Abcam), HIF-1α (1:1000, ab179483, Abcam), Claudin-5 (1:1000, 49564T, CST), Occludin (1:1000, 40-4700, Invitrogen), ZO-1 (1:1000, 21773, Proteintech), and β-actin (1:1500, 66009-1, Proteintech). After incubation with HRP-conjugated secondary antibody against mouse IgG (1:2000, SA00001-1, Proteintech) or rabbit IgG (1:2000, SA00001-2, Proteintech), signals were visualized by ECL (WBKLS0500, Millipore) and quantified using ImageJ.

### Immunofluorescence staining

Frozen brain sections (10 μm) were fixed, permeabilized, blocked with 10% donkey serum/0.5% Triton X-100, and incubated overnight at 4 °C with primary antibodies against Sphk1 (1:50, 10670-1-AP, Proteintech), GFAP (1:2000, PA1-10004, Invitrogen), NeuN (1:500, 94403S, CST), CD31 (1:500, MAB1398Z, Merck), Iba-1 (1:500, ab283319, Abcam), Bsg (1:200, ab188190, Abcam), or MMP-9 (1:100, ab283575, Abcam). After washing, slides were incubated with fluorescent dye-conjugated secondary antibodies (1:500, Jackson Immuno Research) and mounted with DAPI-containing medium. Images were captured using a Zeiss microscope (Zeiss Axio Imager Z2 with Apotome.2) and analyzed with ImageJ.

### Blood-brain barrier permeability assays

BBB integrity was assessed by Sulfo-NHS-Biotin extravasation, IgG immunofluorescence, and HRP staining. For Sulfo-NHS-Biotin assay, mice received a retro-orbital injection of 200 µl Sulfo-NHS-biotin (4 mg per mouse). Following a 30-minutes circulation period, the mice were anesthetized, perfused, and fixed as described above. Subsequently, 10-µm coronal brain sections were prepared and co-stained with anti-CD31 antibody (1:500, MAB1398Z, Merck) and streptavidin-488 (1:2000, S32354, Invitrogen) to visualize all blood vessels and areas of Sulfo-NHS-Biotin extravasation, respectively. For HRP staining, mice received an intravenous injection of HRP (0.5 mg/g body weight, P8375, Sigma-Aldrich) 1-hour before euthanasia. Brains were fixed, sectioned at 50 μm, and reacted with DAB substrate (HY-15912, MCE) to visualize HRP leakage. For IgG staining, sections were incubated with Alexa Fluor 488-conjugated anti-mouse IgG (1:500, Jackson Immuno Research) to detect endogenous IgG extravasation.

### Hematoma volume and brain edema

Hematoma volume was measured on coronal brain sections stained with luxol fast blue (C0631S, Beyotime). Images were captured, and the hematoma area was traced using ImageJ. Total volume was calculated by multiplying the sum of areas by section thickness and interval. Brain edema was evaluated by T2-weighted MRI on a 9.4 T small-animal scanner (uMR930, United Imaging). Brain swelling was quantified as (ipsilateral area - contralateral area)/ipsilateral area × 100%.

### Transmission electron microscopy

Mice were intravenously injected with HRP (0.5 mg/g body weight, P8375, Sigma-Aldrich) for 1-hour and then perfused with fixative containing 2.5% glutaraldehyde and 2% paraformaldehyde. Perihematomal tissue was post-fixed, embedded in epoxy resin, and sectioned at 80 nm. Ultrathin sections were stained with uranyl acetate and lead citrate and examined under a JEM-1400Plus electron microscope (JEOL).

### Flow cytometry and endothelial cell sorting

Peri-hematomal tissues from AAV-treated ICH mice were digested as described above and brain endothelial cells were sorted as described previously (*27*). Single-cell suspensions were stained with antibodies against CD45-PE (1:200, 1075144, BD Biosciences), CD31-PE-Cy7 (1:200, 2410868, Invitrogen), and PDGFRβ-APC (1:200, 2127128, Invitrogen). Live cells (7-AAD⁻), CD45⁻, CD31⁺, PDGFRβ⁻, and GFP⁺ (AAV-infected) endothelial cells were sorted using a Sony MA900 cell sorter. Sorted cells were collected for RNA extraction and transcriptome sequencing.

### RNA sequencing and ATAC-Seq

Total RNA from FACS-sorted endothelial cells was used for library preparation with the Illumina NovaSeq 6000 platform (paired-end 150 bp). Raw reads were filtered, aligned to the mouse genome (GRCm38) with STAR, and quantified using featureCounts. Differential expression analysis was performed with edgeR (FDR < 0.05, |log₂FC| > 1). GO, KEGG, Reactome, and GSEA analyses were conducted using clusterProfiler. Protein-protein interaction networks were visualized with Cytoscape. ATAC-seq was performed on peri-hematomal tissues from ICH mice treated with AAV-shRNA-NC or AAV-shRNA-Sphk1. Nuclei were isolated and tagmented using Tn5 transposase (Illumina). Libraries were sequenced on an Illumina X Plus-25B platform. Reads were aligned to mm10 with Bowtie2, and peaks were called using MACS2. Motif enrichment was analyzed with HOMER.

### Transendothelial electrical resistance (TEER)

Primary mouse brain endothelial cells were seeded on Transwell inserts (0.4 μm pore, Corning) and cultured until confluence. After exposure to hemin/hypoxia with or without PF543, TEER was measured using a Millicell ERS-2 volt-ohm meter (Millipore). Resistance values were normalized to the insert area.

### Dual-luciferase reporter assay

The mouse *Bsg* promoter region (∼2 kb) was cloned into pGL3-Basic (Promega) to generate pGL3-Bsg-pro. HEK293T cells were co-transfected with pGL3-Bsg-pro, pRL-TK (renilla internal control), and either pcDNA3.1-HIF-1α or empty vector using Lipofectamine 3000 (L3000015, Invitrogen). Cells were cultured under hemin/hypoxia conditions for 24-hours and treated with PF543 (8 μM, S7177, Selleck) or CAY10444 (10 μM, HY-119401, MCE) as indicated. Luciferase activities were measured using the Dual-luciferase reporter assay system (E1910, Promega). Firefly luciferase activity was normalized to renilla activity.

## Statistical analysis

Data are presented as mean ± SEM. Comparisons between two groups were performed by unpaired Student’s *t*-test; multiple groups were compared by one-way or two-way ANOVA followed by Tukey’s post hoc test. Statistical significance was set at *p* < 0.05. All analyses were conducted using GraphPad Prism 9.0.

## Data availability

The relevant data for the findings of this study can be obtained from the corresponding author upon reasonable request.

## Acknowledgements

This work was funded with these programs:

1. National Natural Science Foundation of China, 32471043.
2. Leading Project of Henan Province Young Medical Research, LJRC2023010.
3. National Key R&D Program of China, 2023YFE0202200.
4. Guangdong Science and Technology Program, 2025B1212150002.
5. Shenzhen Medical Research Fund, D2403002.
6. National Natural Science Foundation of China, 82503692.

## Competing interests

There are no financial conflicts of interest to disclose.

## Author contributions

Study conception and design: Fuyou Guo, Junlei Chang & Zhihua Li Data collection: Mengzhao Feng, Qi Qin, Kaiyuan Zhang & Min Yu

Analysis and interpretation of results: Mengzhao Feng, Min Yu & Fang Wang Draft manuscript preparation: Mengzhao Feng

Critical revision of the article: Fuyou Guo, Junlei Chang & Zhihua Li

Other (study supervision, fundings, materials, etc…): Fuyou Guo, Junlei Chang & Zhihua Li

All authors reviewed the results and approved the final version of the manuscript.

## Ethical approval

The Ethics Committee for human experiments of Zhengzhou University granted ethical approval for all human sample collection and analysis (Approval number: 2021-KY-0156). All animal experiments were reviewed and approved by the Institutional Animal Care and Use Committee (IACUC) of Shenzhen Institutes of Advanced Technology, Chinese Academy of Sciences (Approval number: SIAT-IACUC-231116-FMZ-A2385).

## Supplementary Materials

Document S1.

Fig. S1. Endothelial cell-specific knockdown of Sphk1 mitigates neurologic impairment at 1-day after ICH in mice.

## Notes

### Competing Interest Statement

The authors have declared no competing interest.

## References

1. L. Ma, X. Hu, L. Song, X. Chen, M. Ouyang, L. Billot, Q. Li, A. Malavera, X. Li, P. Munoz-Venturelli, A. de Silva, N. H. Thang, K. W. Wahab, J. D. Pandian, M. Wasay, O. M. Pontes-Neto, C. Abanto, A. Arauz, H. Shi, G. Tang, S. Zhu, X. She, L. Liu, Y. Sakamoto, S. You, Q. Han, B. Crutzen, E. Cheung, Y. Li, X. Wang, C. Chen, F. Liu, Y. Zhao, H. Li, Y. Liu, Y. Jiang, L. Chen, B. Wu, M. Liu, J. Xu, C. You, C. S. Anderson, I. Investigators, The third Intensive Care Bundle with Blood Pressure Reduction in Acute Cerebral Haemorrhage Trial (INTERACT3): an international, stepped wedge cluster randomised controlled trial. Lancet 402, 27–40 (2023).

2. G. B. D. C. o. D. Collaborators, Global burden of 288 causes of death and life expectancy decomposition in 204 countries and territories and 811 subnational locations, 1990-2021: a systematic analysis for the Global Burden of Disease Study 2021. Lancet 403, 2100–2132 (2024).

3. G. Pradilla, J. J. Ratcliff, A. J. Hall, B. R. Saville, J. W. Allen, G. Paulon, A. McGlothlin, R. J. Lewis, M. Fitzgerald, A. F. Caveney, X. T. Li, M. Bain, J. Gomes, B. Jankowitz, G. Zenonos, B. J. Molyneaux, J. Davies, A. Siddiqui, M. R. Chicoine, S. G. Keyrouz, J. A. Grossberg, M. V. Shah, R. Singh, B. N. Bohnstedt, M. Frankel, D. W. Wright, D. L. Barrow, E. t. investigators, E. T. Investigators, Trial of Early Minimally Invasive Removal of Intracerebral Hemorrhage. N Engl J Med 390, 1277–1289 (2024).

4. S. N. Ohashi, J. H. DeLong, M. G. Kozberg, D. J. Mazur-Hart, S. J. van Veluw, N. J. Alkayed, L. H. Sansing, Role of Inflammatory Processes in Hemorrhagic Stroke. Stroke 54, 605–619 (2023).

5. X. Li, X. Gao, W. Zhang, M. Liu, Z. Han, M. Li, P. Lei, Q. Liu, Microglial replacement in the aged brain restricts neuroinflammation following intracerebral hemorrhage. Cell Death Dis 13, 33 (2022).

6. X. Hu, M. Ouyang, J. Xu, Y. Liu, X. Li, Y. Jiang, X. Chen, L. Billot, M. Q. Li, A. Malavera, P. M. O. Venturelli, A. de Silva, N. H. Thang, K. W. Wahab, J. D. Pandian, M. Wasay, O. M. Pontes-Neto, C. Abanto, A. Arauz, Z. Li, M. Chen, X. Wang, C. Yang, X. Xin, D. Jiang, J. Zheng, Z. Yu, A. Xiao, C. Tao, L. Chen, B. Wu, H. Li, C. S. Anderson, C. You, L. Song, L. Ma, I. Investigators, Surgical outcomes from haematoma evacuation for intracerebral haemorrhage in the INTERACT3 study. Lancet Reg Health West Pac 62, 101669 (2025).

7. S. M. Greenberg, W. C. Ziai, C. Cordonnier, D. Dowlatshahi, B. Francis, J. N. Goldstein, J. C. Hemphill3rd, R. Johnson, K. M. Keigher, W. J. Mack, J. Mocco, E. J. Newton, I. M. Ruff, L. H. Sansing, S. Schulman, M. H. Selim, K. N. Sheth, N. Sprigg, K. S. Sunnerhagen, A. American Heart Association/American Stroke, 2022 Guideline for the Management of Patients With Spontaneous Intracerebral Hemorrhage: A Guideline From the American Heart Association/American Stroke Association. Stroke 53, e282–e361 (2022).

8. A. S. Arthur, B. S. Jahromi, P. S. Saphier, C. M. Nickele, R. W. Ryan, P. Vajkoczy, C. M. Schirmer, C. P. Kellner, C. C. Matouk, E. J. Arias, J. S. Ullman, M. R. Levitt, Z. A. Hage, D. J. Fiorella, M. S. Investigators, Collaborators, Minimally Invasive Surgery vs Medical Management Alone for Intracerebral Hemorrhage: The MIND Randomized Clinical Trial. JAMA Neurol 82, 1113–1121 (2025).

9. H. Zhang, N. Xing, J. Wang, J. Zhang, C. Jia, Y. Li, H. Fan, Y. Liu, F. Dialameh, N. Cheng, Y. Sun, J. Wang, M. Wang, M. Wu, X. Yin, W. Zhu, J. Li, J. Zhang, C. Jiang, F. Xing, M. Zille, X. Fan, X. Chen, J. Wang, Histopathological and ultrastructural changes in different cell types during ischemic and hemorrhagic stroke. Ageing Res Rev 111, 102846 (2025).

10. D. Song, Y. B. Ji, X. W. Huang, Y. Z. Ma, C. Fang, L. H. Qiu, X. X. Tan, Y. M. Chen, S. N. Wang, J. Chang, F. Guo, Lithium attenuates blood-brain barrier damage and brain edema following intracerebral hemorrhage via an endothelial Wnt/beta-catenin signaling-dependent mechanism in mice. CNS Neurosci Ther 28, 862–872 (2022).

11. S. X. Shi, Y. Xiu, Y. Li, M. Yuan, K. Shi, Q. Liu, X. Wang, W. N. Jin, CD4(+) T cells aggravate hemorrhagic brain injury. Sci Adv 9, eabq0712 (2023).

12. O. A. Jones, S. Mohamed, R. Hinz, A. Paterson, O. A. Sobowale, B. R. Dickie, L. M. Parkes, A. R. Parry-Jones, Neuroinflammation and blood-brain barrier breakdown in acute, clinical intracerebral hemorrhage. J Cereb Blood Flow Metab 45, 233–243 (2025).

13. P. Jia, Q. Peng, X. Fan, Y. Zhang, H. Xu, J. Li, H. Sonita, S. Liu, A. Le, Q. Hu, T. Zhao, S. Zhang, J. Wang, M. Zille, C. Jiang, X. Chen, J. Wang, Immune-mediated disruption of the blood-brain barrier after intracerebral hemorrhage: Insights and potential therapeutic targets. CNS Neurosci Ther 30, e14853 (2024).

14. Y. Ji, Q. Gao, Y. Ma, F. Wang, X. Tan, D. Song, R. L. C. Hoo, Z. Wang, X. Ge, H. Han, F. Guo, J. Chang, An MMP-9 exclusive neutralizing antibody attenuates blood-brain barrier breakdown in mice with stroke and reduces stroke patient-derived MMP-9 activity. Pharmacol Res 190, 106720 (2023).

15. C. P. Profaci, R. N. Munji, R. S. Pulido, R. Daneman, The blood-brain barrier in health and disease: Important unanswered questions. J Exp Med 217, (2020).

16. R. N. Munji, A. L. Soung, G. A. Weiner, F. Sohet, B. D. Semple, A. Trivedi, K. Gimlin, M. Kotoda, M. Korai, S. Aydin, A. Batugal, A. C. Cabangcala, P. G. Schupp, M. C. Oldham, T. Hashimoto, L. J. Noble-Haeusslein, R. Daneman, Profiling the mouse brain endothelial transcriptome in health and disease models reveals a core blood-brain barrier dysfunction module. Nat Neurosci 22, 1892–1902 (2019).

17. L. Li, H. Wang, J. Zhang, Y. Sha, F. Wu, S. Wen, L. He, L. Sheng, Q. You, M. Shi, L. Liu, H. Zhou, SPHK1 deficiency protects mice from acetaminophen-induced ER stress and mitochondrial permeability transition. Cell Death Differ 27, 1924–1937 (2020).

18. X. Ji, Z. Chen, Q. Wang, B. Li, Y. Wei, Y. Li, J. Lin, W. Cheng, Y. Guo, S. Wu, L. Mao, Y. Xiang, T. Lan, S. Gu, M. Wei, J. Z. Zhang, L. Jiang, J. Wang, J. Xu, N. Cao, Sphingolipid metabolism controls mammalian heart regeneration. Cell Metab 36, 839–856 e838 (2024).

19. M. Feng, Q. Qin, K. Zhang, F. Wang, D. Song, M. Li, Y. An, Z. Li, F. Guo, Sphk2 in ischemic stroke pathogenesis: Roles, mechanisms, and regulation strategies. Ageing Res Rev 111, 102844 (2025).

20. P. Zhou, L. Zhou, Y. Shi, Z. Li, L. Liu, L. Zuo, J. Zhang, S. Liang, J. Kang, S. Du, J. Yang, Z. Sun, X. Zhang, Neuroprotective Effects of Danshen Chuanxiongqin Injection Against Ischemic Stroke: Metabolomic Insights by UHPLC-Q-Orbitrap HRMS Analysis. Front Mol Biosci 8, 630291 (2021).

21. J. Xie, T. Zhang, P. Li, D. Wang, T. Liu, S. Xu, Dihydromyricetin Attenuates Cerebral Ischemia Reperfusion Injury by Inhibiting SPHK1/mTOR Signaling and Targeting Ferroptosis. Drug Des Devel Ther 16, 3071–3085 (2022).

22. D. Su, Y. Cheng, S. Li, D. Dai, W. Zhang, M. Lv, Sphk1 mediates neuroinflammation and neuronal injury via TRAF2/NF-kappaB pathways in activated microglia in cerebral ischemia reperfusion. J Neuroimmunol 305, 35–41 (2017).

23. W. Liu, X. Zhou, K. Zeng, C. Nie, J. Huang, L. Zhu, D. Pei, Y. Zhang, Study on the action mechanism of Buyang Huanwu Decoction against ischemic stroke based on S1P/S1PR1/PI3K/Akt signaling pathway. J Ethnopharmacol 312, 116471 (2023).

24. B. P. Gaire, J. W. Choi, Sphingosine 1-Phosphate Receptors in Cerebral Ischemia. Neuromolecular Med 23, 211–223 (2021).

25. D. Cong, Y. Yu, Y. Meng, X. Qi, Dexmedetomidine (Dex) exerts protective effects on rat neuronal cells injured by cerebral ischemia/reperfusion via regulating the Sphk1/S1P signaling pathway. J Stroke Cerebrovasc Dis 32, 106896 (2023).

26. M. Feng, Y. An, Q. Qin, I. H. Fong, K. Zhang, F. Wang, D. Song, M. Li, M. Yu, C. T. Yeh, J. Chang, F. Guo, Sphk1/S1P pathway promotes blood-brain barrier breakdown after intracerebral hemorrhage through inducing Nlrp3-mediated endothelial cell pyroptosis. Cell Death Dis 15, 926 (2024).

27. M. Yu, Y. Nie, J. Yang, S. Yang, R. Li, V. Rao, X. Hu, C. Fang, S. Li, D. Song, F. Guo, M. P. Snyder, H. Y. Chang, C. J. Kuo, J. Xu, J. Chang, Integrative multi-omic profiling of adult mouse brain endothelial cells and potential implications in Alzheimer’s disease. Cell Rep 42, 113392 (2023).

28. Z. Shi, J. Li, Z. Feng, C. Fang, Y. Wang, L. Qiu, J. Liu, F. Wang, Z. N. Guo, Y. Yang, K. Huang, J. Chang, Y. Ma, Dickkopf-related protein 2 impairs neurovascular Wnt signalling and worsens stroke outcome. Eur Heart J, (2025).

29. X. Huang, P. Wei, C. Fang, M. Yu, S. Yang, L. Qiu, Y. Wang, A. Xu, R. L. C. Hoo, J. Chang, Compromised endothelial Wnt/beta-catenin signaling mediates the blood-brain barrier disruption and leads to neuroinflammation in endotoxemia. J Neuroinflammation 21, 265 (2024).

30. Q. He, Y. Ma, C. Fang, Z. Deng, F. Wang, Y. Qu, M. Yin, R. Zhao, D. Zhang, F. Guo, Y. Yang, J. Chang, Z. N. Guo, Remote ischemic conditioning attenuates blood-brain barrier disruption after recombinant tissue plasminogen activator treatment via reducing PDGF-CC. Pharmacol Res 187, 106641 (2023).

31. Z. Feng, C. Fang, Y. Ma, J. Chang, Obesity-induced blood-brain barrier dysfunction: phenotypes and mechanisms. J Neuroinflammation 21, 110 (2024).

32. J. Chang, M. R. Mancuso, C. Maier, X. Liang, K. Yuki, L. Yang, J. W. Kwong, J. Wang, V. Rao, M. Vallon, C. Kosinski, J. J. Zhang, A. T. Mah, L. Xu, L. Li, S. Gholamin, T. F. Reyes, R. Li, F. Kuhnert, X. Han, J. Yuan, S. H. Chiou, A. D. Brettman, L. Daly, D. C. Corney, S. H. Cheshier, L. D. Shortliffe, X. Wu, M. Snyder, P. Chan, R. G. Giffard, H. Y. Chang, K. Andreasson, C. J. Kuo, Gpr124 is essential for blood-brain barrier integrity in central nervous system disease. Nat Med 23, 450–460 (2017).

33. C. P. Profaci, S. S. Harvey, K. Bajc, T. Z. Zhang, D. A. Jeffrey, A. Z. Zhang, K. M. Nemec, H. Davtyan, C. A. O’Brien, G. L. McKinsey, A. Longworth, T. P. McMullen, J. K. Capocchi, J. G. Gonzalez, D. A. Lawson, T. D. Arnold, D. Davalos, M. Blurton-Jones, F. Dabertrand, F. C. Bennett, R. Daneman, Microglia are not necessary for maintenance of blood-brain barrier properties in health, but PLX5622 alters brain endothelial cholesterol metabolism. Neuron 112, 2910–2921 e2917 (2024).

34. M. A. Petersen, J. K. Ryu, K. J. Chang, A. Etxeberria, S. Bardehle, A. S. Mendiola, W. Kamau-Devers, S. P. J. Fancy, A. Thor, E. A. Bushong, B. Baeza-Raja, C. A. Syme, M. D. Wu, P. E. Rios Coronado, A. Meyer-Franke, S. Yahn, L. Pous, J. K. Lee, C. Schachtrup, H. Lassmann, E. J. Huang, M. H. Han, M. Absinta, D. S. Reich, M. H. Ellisman, D. H. Rowitch, J. R. Chan, K. Akassoglou, Fibrinogen Activates BMP Signaling in Oligodendrocyte Progenitor Cells and Inhibits Remyelination after Vascular Damage. Neuron 96, 1003–1012 e1007 (2017).

35. J. Niu, H. H. Tsai, K. K. Hoi, N. Huang, G. Yu, K. Kim, S. E. Baranzini, L. Xiao, J. R. Chan, S. P. J. Fancy, Aberrant oligodendroglial-vascular interactions disrupt the blood-brain barrier, triggering CNS inflammation. Nat Neurosci 22, 709–718 (2019).

36. M. Martin, S. Vermeiren, N. Bostaille, M. Eubelen, D. Spitzer, M. Vermeersch, C. P. Profaci, E. Pozuelo, X. Toussay, J. Raman-Nair, P. Tebabi, M. America, A. De Groote, L. E. Sanderson, P. Cabochette, R. F. V. Germano, D. Torres, S. Boutry, A. de Kerchove d’Exaerde, E. J. Bellefroid, T. N. Phoenix, K. Devraj, B. Lacoste, R. Daneman, S. Liebner, B. Vanhollebeke, Engineered Wnt ligands enable blood-brain barrier repair in neurological disorders. Science 375, eabm4459 (2022).

37. J. H. Lawrence, A. Patel, M. W. King, C. J. Nadarajah, R. Daneman, E. S. Musiek, Microglia drive diurnal variation in susceptibility to inflammatory blood-brain barrier breakdown. JCI Insight 9, (2024).

38. M. Blanchette, K. Bajc, B. D. Gastfriend, C. P. Profaci, N. Ruderisch, C. E. Dorrier, G. Zhong, R. Cuevas-Diaz Duran, S. S. Harvey, I. H. Garcia-Pak, L. Pintaric, M. Leclerc, L. Reveret, V. Emond, A. Wang, D. Pant, L. T. Tsai, F. Calon, N. Isoherranen, S. P. Palecek, E. V. Shusta, J. Wu, R. Daneman, Regional heterogeneity of the blood-brain barrier. Nat Commun 16, 7332 (2025).

39. W. Sheng, Z. Wu, J. Wei, J. Wang, S. Zhang, Z. Ding, J. Zhong, D. Deng, Z. Zhong, Y. Yin, Y. Li, Q. Wang, Astrocyte-derived CXCL10 exacerbates endothelial cells pyroptosis and blood-brain barrier disruption via CXCR3/cGAS/AIM2 pathway after intracerebral hemorrhage. Cell Death Discov 11, 373 (2025).

40. J. Lin, Y. Xu, P. Guo, Y. J. Chen, J. Zhou, M. Xia, B. Tan, X. Liu, H. Feng, Y. Chen, CCL5/CCR5-mediated peripheral inflammation exacerbates blood‒brain barrier disruption after intracerebral hemorrhage in mice. J Transl Med 21, 196 (2023).

41. X. Lei, E. Sun, X. Ru, Y. Quan, X. Chen, Q. Zhang, Y. Lu, Q. Huang, Y. Chen, W. Li, H. Feng, Y. Yang, R. Hu, Acetylation of alpha-tubulin restores endothelial cell injury and blood-brain barrier disruption after intracerebral hemorrhage in mice. Exp Mol Med 57, 1064–1077 (2025).

42. Q. Qin, M. Feng, K. Zhang, Z. Mo, Y. Liu, Y. Ma, X. Liu, Basigin in cerebrovascular diseases: Roles, mechanisms, and therapeutic target potential. Eur J Pharmacol 989, 177232 (2025).

43. Y. Liu, Z. Li, S. Khan, R. Zhang, R. Wei, Y. Zhang, M. Xue, V. W. Yong, Neuroprotection of minocycline by inhibition of extracellular matrix metalloproteinase inducer expression following intracerebral hemorrhage in mice. Neurosci Lett 764, 136297 (2021).

44. Y. Liu, Q. Bai, V. W. Yong, M. Xue, EMMPRIN Promotes the Expression of MMP-9 and Exacerbates Neurological Dysfunction in a Mouse Model of Intracerebral Hemorrhage. Neurochem Res 47, 2383–2395 (2022).

45. J. Zhuang, Q. Shang, F. Rastinejad, D. Wu, Decoding Allosteric Control in Hypoxia-Inducible Factors. J Mol Biol 436, 168352 (2024).

46. G. L. Wang, B. H. Jiang, E. A. Rue, G. L. Semenza, Hypoxia-inducible factor 1 is a basic-helix-loop-helix-PAS heterodimer regulated by cellular O2 tension. Proc Natl Acad Sci U S A 92, 5510–5514 (1995).

47. G. L. Semenza, F. Agani, G. Booth, J. Forsythe, N. Iyer, B. H. Jiang, S. Leung, R. Roe, C. Wiener, A. Yu, Structural and functional analysis of hypoxia-inducible factor 1. Kidney Int 51, 553–555 (1997).

48. M. D. Michaud, G. A. Robitaille, J. P. Gratton, D. E. Richard, Sphingosine-1-phosphate: a novel nonhypoxic activator of hypoxia-inducible factor-1 in vascular cells. Arterioscler Thromb Vasc Biol 29, 902–908 (2009).

49. X. Ke, F. Fei, Y. Chen, L. Xu, Z. Zhang, Q. Huang, H. Zhang, H. Yang, Z. Chen, J. Xing, Hypoxia upregulates CD147 through a combined effect of HIF-1alpha and Sp1 to promote glycolysis and tumor progression in epithelial solid tumors. Carcinogenesis 33, 1598–1607 (2012).

50. D. Xu, Q. Gao, F. Wang, Q. Peng, G. Wang, Q. Wei, S. Lei, S. Zhao, L. Zhang, F. Guo, Sphingosine-1-phosphate receptor 3 is implicated in BBB injury via the CCL2-CCR2 axis following acute intracerebral hemorrhage. CNS Neurosci Ther 27, 674–686 (2021).

51. T. Krolak, K. Y. Chan, L. Kaplan, Q. Huang, J. Wu, Q. Zheng, V. Kozareva, T. Beddow, I. G. Tobey, S. Pacouret, A. T. Chen, Y. A. Chan, D. Ryvkin, C. Gu, B. E. Deverman, A High-Efficiency AAV for Endothelial Cell Transduction Throughout the Central Nervous System. Nat Cardiovasc Res 1, 389–400 (2022).

52. Y. Wang, M. Nakayama, M. E. Pitulescu, T. S. Schmidt, M. L. Bochenek, A. Sakakibara, S. Adams, A. Davy, U. Deutsch, U. Luthi, A. Barberis, L. E. Benjamin, T. Makinen, C. D. Nobes, R. H. Adams, Ephrin-B2 controls VEGF-induced angiogenesis and lymphangiogenesis. Nature 465, 483–486 (2010).

